# Treatment of a metabolic liver disease by *in vivo* prime editing in mice

**DOI:** 10.1101/2021.08.17.456632

**Authors:** Desirée Böck, Tanja Rothgangl, Lukas Villiger, Lukas Schmidheini, Nicholas Mathis, Eleonora Ioannidi, Susanne Kreutzer, Zacharias Kontarakis, Nicole Rimann, Hiu Man Grisch-Chan, Beat Thöny, Gerald Schwank

**Affiliations:** Institute of Pharmacology and Toxicology, University of Zurich, Zurich, Switzerland; Institute of Molecular Health Sciences, ETH Zurich, Zurich, Switzerland; Functional Genomics Center Zurich, ETH Zurich/University of Zurich, Zurich, Switzerland; Genome Engineering and Measurement Laboratory (GEML), ETH Zurich, Zurich, Switzerland; Division of Metabolism and Children’s Research Center, University Children’s Hospital Zurich, Zurich, Switzerland; Zurich Center for Integrative Human Physiology, Zurich, Switzerland; Neuroscience Center Zurich, Zurich, Switzerland

## Abstract

Prime editing is a highly versatile CRISPR-based genome editing technology with the potential to correct the vast majority of pathogenic mutations *(1)*. However, correction of a disease phenotype *in vivo* in somatic tissues has not been demonstrated thus far. Here, we establish proof-of-concept for *in vivo* prime editing and repair the metabolic liver disease phenylketonuria (PKU) in mice. We first developed a size-reduced *Sp*Cas9 prime editor (PE) lacking the RNaseH domain of the reverse transcriptase (PE2^Δ*RnH*^), and a linker- and NLS-optimized intein-split PE construct (PE2 p.1153) for delivery by adeno-associated virus (AAV) vectors. Systemic dual AAV-mediated delivery of this variant into the liver of neonatal mice enabled installation of a transversion mutation at the *Dnmt1* locus with an average efficiency of 15%, and delivery of unsplit PE2^Δ*RnH*^ using human adenoviral vector 5 (AdV5) further increased editing rates to 58%. PE2^Δ*RnH*^-encoding AdV5 was also used to correct the disease-causing mutation of the phenylalanine hydroxylase *(Pah)*^*enu2*^ allele in phenylketonuria (PKU) mice with an average efficiency of 8% (up to 17.3%), leading to therapeutic reduction of blood phenylalanine (L-Phe) levels. Our study demonstrates *in vivo* prime editing in the liver with high precision and editing rates sufficient to treat a number of metabolic liver diseases, emphasizing the potential of prime editing for future therapeutic applications.

**One Sentence Summary:** *In vivo* prime editing corrects phenylketonuria in mice.

## INTRODUCTION

More than 75’000 disease-associated genetic variants have been identified to date. The majority of them are caused by small insertion/deletion mutations or base substitutions *(2)*. Programmable CRISPR-Cas9 nucleases have previously been applied to correct pathogenic alleles *in vivo (3, 4)*. However, these systems rely on homology-directed repair (HDR) mechanisms to install precise edits *(5–7)*, making them inefficient and error-prone, in particular in somatic tissues *(8, 9)*. Base editors (BEs) are CRISPR-based genome editing tools where nuclease-impaired Cas9 directs a deaminase to the locus of interest in order to install single nucleotide conversions. They enable precise genome editing independent of HDR *(10, 11)*, and also operate with high efficiency and accuracy in somatic tissues *in vivo (12–18)*. However, BEs can only install transition substitutions (A-T to G-C or C-T to T-A conversions), and are therefore not applicable to a large proportion of known pathogenic mutations *(2)*.

Prime editors (PEs) are more recently developed genome editing tools, which consist of an RNA-programmable *Sp*Cas9 nickase (H840A) fused to an engineered reverse transcriptase (RT). Additional to a guide-scaffold sequence, the prime editing guide RNA (pegRNA) contains a primer binding site (PBS) and a template sequence, which is reverse transcribed into DNA and copied into the targeted locus. Since PEs directly write new genetic information, they are extremely versatile, and allow the installation of all possible nucleotide conversions as well as short deletions and insertions up to 80 bp *(1)*. To improve editing efficiency of PEs, mutations that enhance the activity of the RT have been introduced (PE2) and a second sgRNA can be designed to nick the non-edited DNA strand either in parallel (PE3) or after installing the edit (PE3b), facilitating DNA repair of the edited DNA strand *(1, 19, 20)*. While prime editing has already been demonstrated *in vitro* in cultured cells *(1, 21–23)* and *in vivo* in mice and *Drosophila (19, 20, 24, 25)*, treatment of animal models for human diseases via prime editing has not been reported yet. Here, we demonstrate *in vivo* prime editing in proliferating hepatocytes of neonatal mice and non-proliferating hepatocytes of adult mice. We report that high vector doses are crucial to achieve efficient editing, and in a proof-of-concept describe the correction of the *Pah*^*enu2*^ mutation with therapeutic reduction of L-Phe levels in PKU mice.

## RESULTS

### Subhead 1: Establishment of the size-reduced PE variant PE2^Δ*RnH*^

*In vivo* prime editing holds great potential for therapeutic applications. However, the large size of PEs (~6.6 kb; Fig. 1a) poses a challenge for *in vivo* delivery via viral and non-viral vectors.

**Figure 1:**
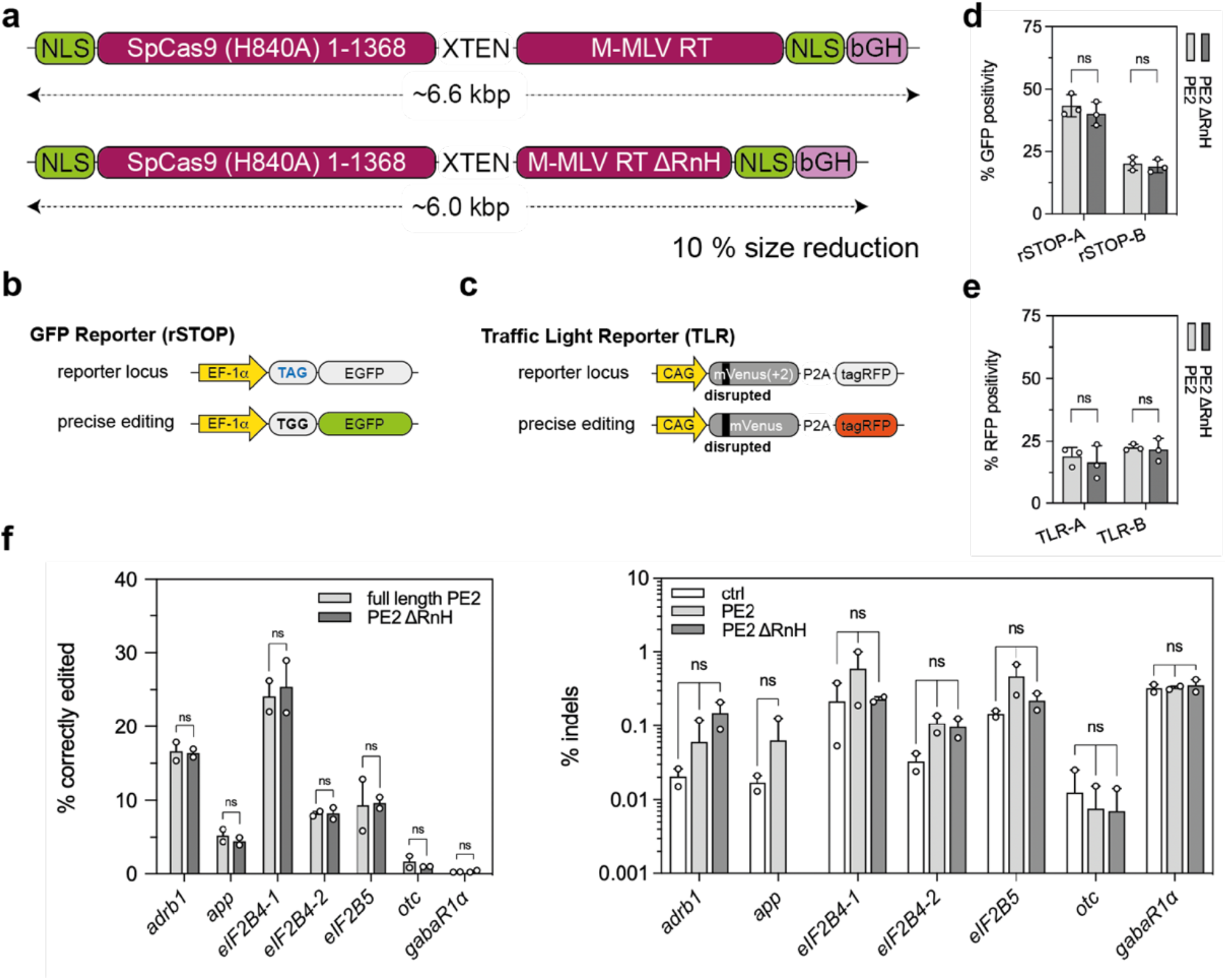
Establishment of the size-reduced PE variant PE^Δ*RnH*^. (**a**) Schematic representation of the full-length PE2 and our size-reduced variant PE2^Δ*RnH*^, lacking the RNaseH (RnH) domain (Δ486-677) of the RT. (**b**) rSTOP reporter: conversion of a TAG stop codon results in GFP expression. (**c**) TLR reporter: correction of a 2 bp frameshift results in tagRFP expression. (**d, e**) Performance of PE2 and PE2^Δ*RnH*^ in rSTOP (d) and TLR (e) reporter cells using 2 different protospacers (labelled as A and B). Editing efficiency is scored by flow cytometry. (**f**) Comparative analyses of on-target editing efficiency and indel formation of PE2 and PE2^Δ*RnH*^ at seven genomic sites (ns, not significant; left panel: two-tailed student’s *t*-test; right panel: two-way ANOVA with Tukey’s multiple comparisons test). pegRNA plasmids were transfected as negative controls. Data from all experiments are represented as mean ± s.d. (three independent experiments; d, e) or range (two independent experiments; f). PE, prime editor; M-MLV, Moloney murine leukemia virus; RT, reverse transcriptase; NLS, nuclear localization signal; bGH, bovine growth hormone polyadenylation signal; EF-1α, eukaryotic translation elongation factor 1α; rSTOP, remove stop codon; adrb1, β1-adrenergic receptor; app, amyloid β-precursor protein; eIF2B, eukaryotic translation initiation factor 2B; otc, ornithine carbamoyltransferase; gabaR1α, gamma-aminobutyric acid receptor subunit α-1.

Since previous studies have indicated that the polymerase and RNaseH domains of the M-MLV RT can function independently of each other *(26, 27)*, we speculated that the RNaseH domain (~0.6 kb), which is required to degrade DNA-RNA heteroduplexes *(28)*, could be negligible in the context of prime editing. Therefore, we compared editing efficiencies of PE2^Δ*RnH*^ (RT^Δ486-677^) and PE2 in two different HEK293T fluorescent reporter cell lines: the rSTOP reporter, in which GFP expression is abolished by a premature TAG stop codon (Fig. 1b), and the traffic light reporter (TLR), where translation of a functional tagRFP is prevented by a 2 bp frameshift (Fig. 1c). Using multiple pegRNAs, we observed comparable editing efficiencies between PE2 and PE2^Δ*RnH*^ by FACS analysis (Fig. 1d, e; Extended data fig. 1). To corroborate these results, we further targeted seven pathogenic mutations (transversions and transitions) at different loci, and analyzed them by deep sequencing. Again, no differences in editing rates at the targeted nucleotide were found between PE2 and PE2^Δ*RnH*^ (Fig. 1f). In addition, we did not observe significant indel formation above background in any of these sites upon PE2 and PE2^Δ*RnH*^ treatment (Fig. 1f). Our data therefore demonstrate that the size-reduced PE2^Δ*RnH*^ harbors comparable efficiency and accuracy as PE2.

### Subhead 2: Establishment of an intein-split PE variant for dual AAV-mediated delivery

Due to their low immunogenicity and broad range of serotype specificity, AAVs are promising candidates for *in vivo* delivery of genome editing tools *(29)*. To stay within the limited AAV packaging capacity (4.7 kb) *(30)*, the coding sequence of Cas9 nucleases or BEs has previously been split onto two separate AAVs using the intein-mediated protein trans-splicing system from cyanobacterium *Nostoc punctiforme* (*Npu*) *(13, 14, 31, 32)*. To optimize the intein-split approach for PE delivery, we assessed the activity of *SpCas9*-PE variants, which were split at different sites. Within the region where both generated PE segments would not exceed the packaging limit of an AAV we identified eight surface-exposed positions with either a Cys, Ser, or Thr at the N-terminal position of the C-intein PE moiety *(33)*. When tested with the rSTOP reporter, previously used split variants (p.573 and p.714) *(14, 31, 32)* showed a substantial reduction in editing activity compared to unsplit PE2 (Fig. 2b), and were outperformed by the PE2-p.1153 variant that maintained 75% of the activity of unsplit PE2 (Fig. 2b). Further analysis of the intein-split architecture of PE2-p.1153 revealed the importance of flexible Glycine-Serine ([GGGGS]3)-linkers between the intein domains and PE segments (Extended data fig. 2a), and showed that removal of the nuclear localization signal (NLS) from the N-intein splice-donor is beneficial for activity (Fig. 2c; Extended data fig. 2b). Notably, we also evaluated potential split sites for circular permutant *Sp*Cas9-PE (Extended data fig. 2c), and for PE variants based on orthogonal Cas9 from *Staphylococcus aureus* (*Sa*Cas), *Staphylococcus auricularis* (*Sauri*Cas), and *Campylobacter jejuni* (*Cj*Cas) (Extended data fig. 3). However, none of the tested full-length variants led to editing rates comparable to *Sp*Cas9-PE, prompting us to continue with the linker- and NLS-optimized PE2-p.1153 variant for *in vivo* experiments.

**Figure 2:**
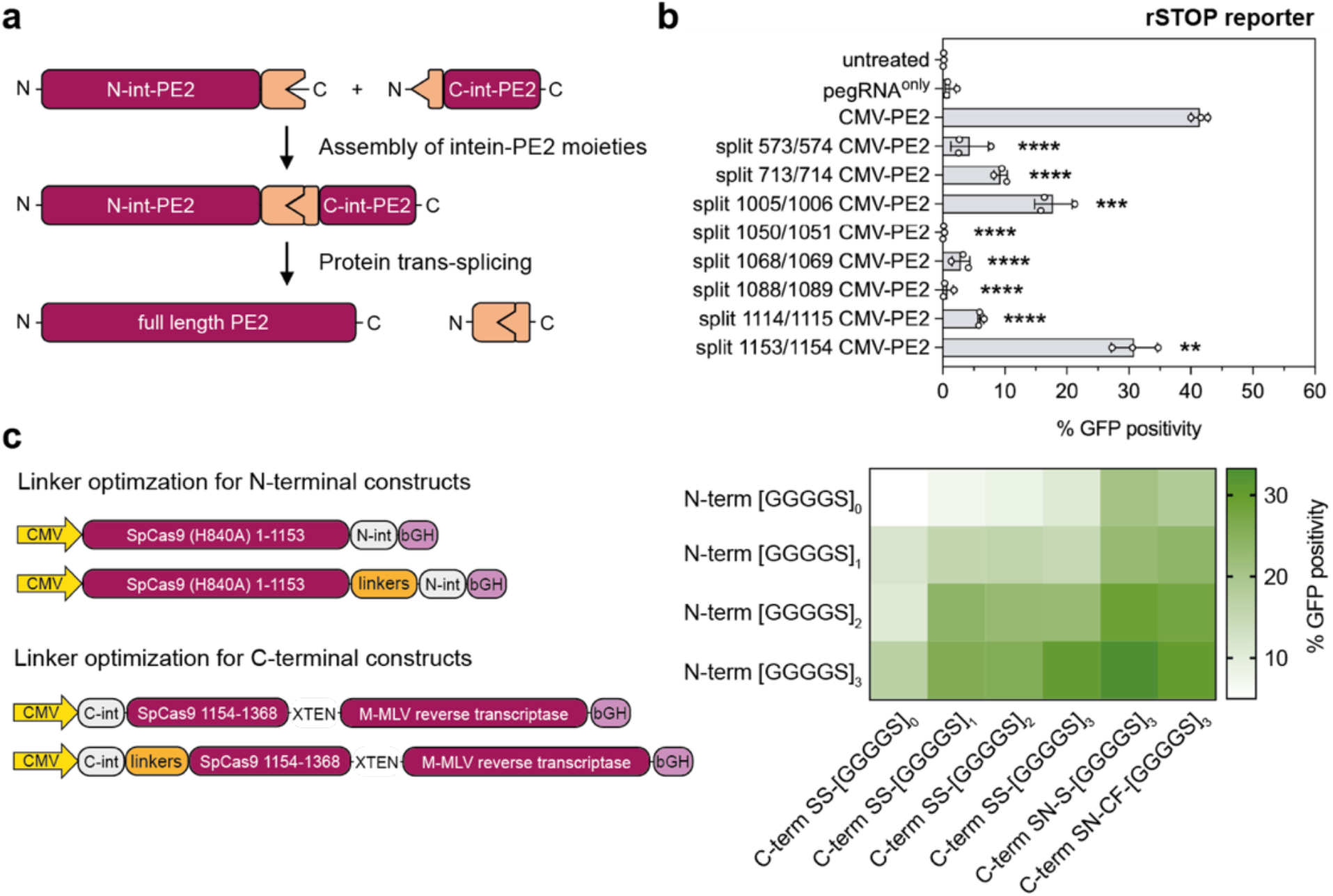
Establishment of an intein-split PE variant for dual AAV-mediated delivery. (**a**) Depiction of the two intein-split PE moieties (N-int-PE2 and C-int-PE2), forming the full-length PE after protein trans-splicing. (**b**) Editing efficiencies of various intein-split PEs with Glycine-Serine ([GGGGS]3)-linkers in the rSTOP reporter. Untreated reporter cells or transfected pegRNA plasmid were used as controls. Efficiencies of intein-split PEs were compared to full-length CMV-PE2 and analyzed using a two-tailed student’s t-test with Welch’s correction. (**c**) Optimization of the linker length (1-3 repeats of the Glycine-Serine linker) and the intein-split splice-acceptor site (amino acid combinations SS, SN-S, or SN-CF) at position p.1153. Left panel: Schematic maps of N- and C-terminal intein-split PE halves, highlighting the linker position. Right panel: Comparison of editing performance of different combinations of optimized N- and C-int-PE2 in the rSTOP GFP reporter. Data of all experiments are depicted as means of at least three independent biological replicates. N-int, N-intein; C-int, C-intein; CMV, human cytomegalovirus promoter; N-term, N-terminal PE half; C-term, C-terminal PE half. *\*\*P>0.005*, *\*\*\*P>0.0005*, *\*\*\*\*P>0.0001*.

### Subhead 3: AAV-mediated prime editing at the *Dnmt1* locus in the mouse liver

To establish *in vivo* prime editing approaches in the liver we employed a pegRNA that has previously been demonstrated to install a G-to-C transversion mutation in the *Dnmt1* gene with high efficiency in mouse embryos*(19)*. We first confirmed efficient on-target editing without significant induction of indels at the endogenous *Dnmt1* locus when co-expressing the pegRNA with PE2^Δ*RnH*^ *in vitro* in Hepa1-6 cells (Extended data fig. 4a). We next generated AAV constructs where intein-split PE2^Δ*RnH*^ and the pegRNA are expressed under the synthetic liver-specific P3 *(34)* and U6 promoter, respectively, and packaged them into AAV2 serotype 8 particles (Fig. 3a; hereafter referred to as AAV8). AAV particles where then systemically delivered into C57BL/6J pups on postnatal day 1 (P1) via the temporal vein at a dose of 5×10^11^ vector genomes (vg) per construct and animal (Fig. 3a, Extended data fig. 4b), and primary hepatocytes were isolated after 4 weeks to analyze editing rates by targeted amplicon deep sequencing. The intended G-to-C conversion was installed with an efficiency of 14.4 ± 6.6%, with no indels above background being detected at the target locus (Fig. 3b). Notably, co-expression of a second sgRNA, which binds to the target locus upon introduction of the novel PAM generated by the G-to-C edit and which was designed by Aida *et al*. with the intention to induce a nick in the unedited strand *(19)*, did not further increase editing rates *in vitro* and *in vivo* (12.2 ± 1.5% and 15.9 ± 7.3%, Extended data fig. 4a, 4b). This is likely due to the fact that unlike for the PE3b approach *(1)*, where the unedited strand is directly nicked after the initial edit is installed, introduction of a novel PAM site requires copying of the edit to the opposite DNA strand before the Cas9 complex can bind. We next analyzed AAV treated mice 12 weeks post injection instead of 4 weeks post injection. However, editing rates did not significantly increase (Extended data fig. 4c), suggesting that AAV-mediated *in vivo* prime editing at the *Dnmt1* locus already reached its plateau at 4 weeks after injection in neonates.

**Figure 3:**
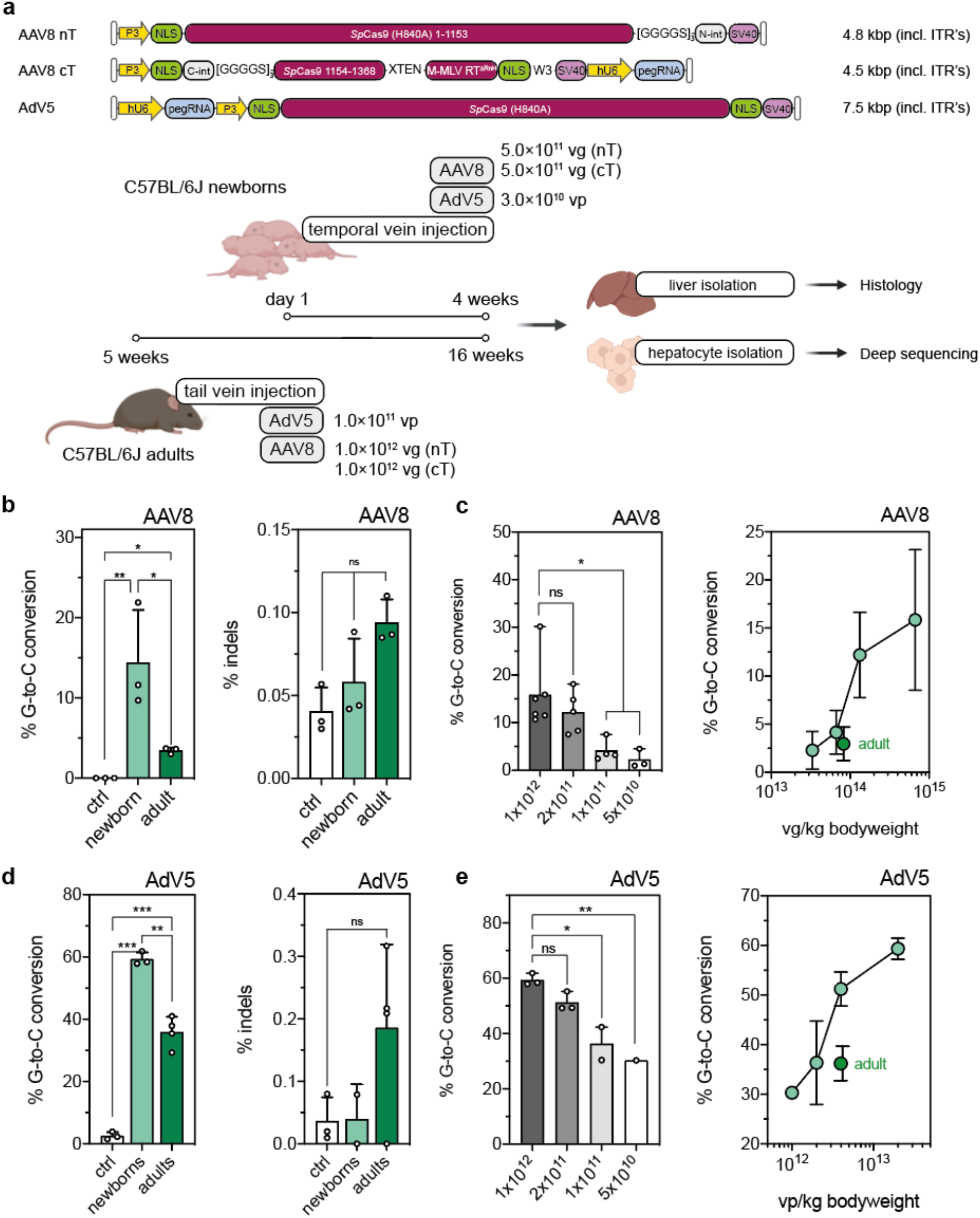
AAV8- and AdV5-mediated prime editing at the *Dnmt1* locus in the mouse liver. **(a)** Schematic outline of the experimental setup with AAV8- or AdV5-mediated prime editing in newborn and adult mice. Constructs used for *in vivo* prime editing at the *Dnmt1* locus in the mouse liver are not depicted to scale. (**b**) Correction and indel rates in newborn and adult animals after AAV8-mediated delivery (injected dose per neonate and adult: 1×10^12^ and 2×10^12^ vg). Untreated mice were used as negative controls. Percentage of sequencing reads with indels around the protospacer region were determined by deep sequencing. (**c**) Editing rates in neonates relative to AAV doses per animal (left panel) and per kg bodyweight (right panel). The respective values for adult mice were added (dark green) for comparison. (**d**) Correction and indel rates in newborn and adult mice at 4 weeks after AdV5-mediated delivery (injected dose per neonate and adult: 3×10^10^ and 1×10^11^ viral particles, vp). (**e**) Editing rates in neonates relative to AdV5 doses per animal (left panel) and per kg bodyweight (right panel). The respective values for adult mice were added (dark green) for comparison. Untreated mice were used as negative controls. Data are represented as mean ± s.d. (n=3-6 mice per group) and were analyzed using a two-way ANOVA with Tukey’s multiple comparisons test (*\*P>0.05; **P<0.005*).

To next assess editing efficiencies in adult animals, we injected 5 week-old C57BL/6J mice via the tail vein at a dose of 1×10^12^ vg per construct and animal. After 12 weeks, hepatocytes were isolated and on-target editing rates were analyzed by deep sequencing. Again, we observed editing of the target base without indel formation above background (Fig. 3b). However, editing rates were significantly lower compared to neonatal animals, with the intended G-to-C conversion being present in 3.4 ± 0.4% of *Dnmt1* alleles (Fig. 3b). One potential explanation for lower editing rates in adult animals is the lack of hepatocyte proliferation compared to neonates. We therefore next induced ectopic hepatocyte proliferation at three or six weeks after PE administration via two-third partial hepatectomy (PHx, Extended data fig. 5a). Nevertheless, proliferating hepatocytes after PHx (confirmed in control mice 3 days after surgery by immunofluorescence, Extended data fig. 5b) did not further increase editing rates (Extended data fig. 5c). An alternative explanation for our observation of reduced editing rates in adult animals is different dosing, as neonates were injected with higher viral doses per kg bodyweight than adults. Indeed, when we injected P1 pups with a 5-, 10-, or 20-fold lower dose of AAV8 we observed a dose-dependent decline of editing rates (Fig. 3c), with neonates receiving a viral dose per kg bodyweight comparable to adults displaying similar C-to-G conversion rates (Fig. 3c).

### Subhead 4: AdV5-mediated prime editing at the *Dnmt1* locus in the mouse liver

Due to the relatively low reconstitution efficiency of full-length proteins from intein-split moieties *(13)* (Extended data fig. 6), and the observed dose-dependent increase in editing rates at the *Dnmt1* locus (Fig. 3c), we hypothesized that delivery of unsplit PEs using a vector with higher packaging capacity could enhance prime editing rates. We therefore delivered a human Adenovirus 5 (AdV5) vector targeting the *Dnmt1* locus with unsplit PE2^Δ*RnH*^ into C57BL/6J adult or neonatal mice (Fig. 3a). In line with our assumption, we observed a substantial increase in editing compared to AAV-treated animals, with rates of 58.2 ± 0.4% and 35.9 ± 4.9% in neonates and adults, respectively (Fig. 3d). Editing efficiencies were also strongly dependent on the AdV5 dose, and declined when a 5-, 10-, or 20-fold lower dose was injected in neonates (Fig. 3e). Notably, despite the high on-target editing efficiency after AdV5-mediated PE2^Δ*RnH*^ delivery indel mutations were still within the range of untreated control animals (Fig. 3d), confirming high precision of *in vivo* prime editing.

### Subhead 5: Establishment of prime editing strategies to correct the *Pah*^*enu2*^ allele

For a number of genetic liver diseases, including PKU, correction rates below 10% could already lead to significant therapeutic effects *(35)*. PKU is an autosomal recessive metabolic liver disease, which is caused by mutations in the phenylalanine hydroxylase (*Pah*) gene. These result in a lowering of functional PAH enzyme levels, leading to toxic accumulation of phenylalanine (L-Phe) and its byproducts in the blood. The *Pah*^*enu2*^ mouse model for PKU carries a homozygous point mutation on exon 7 (c.835T>C; p.F263S), resulting in abnormally high blood L-Phe levels of ~2000 μmol/L *(36)*. While correction of the mutation using cytidine BEs has led to a reduction in blood L-Phe levels below the therapeutic threshold and full restoration of the PKU phenotype *(13, 16)*, product purity of the edited locus was relatively low, with more than half of the edited alleles containing either bystander or indel mutations *(13)*.

To assess whether the *Pah*^*enu2*^ locus could be corrected with higher precision using prime editing, we first evaluated on-target editing rates and indel formation using various pegRNAs *in vitro* in a HEK293T cell line with stably integrated exon 7 of the *Pah*^*enu2*^ allele. We tested pegRNAs for two different *Sp*Cas9 protospacers and for one *Sp*RY protospacer, which could also be targeted with a PE3b sgRNA (Fig 4a, 4b, Extended data fig. 7a). pegRNAs harbored a 13-nucleotide long PBS domain combined with 16- (mPKU-*.1) or 19- (mPKU-*.2) nucleotide long RT templates. Deep amplicon sequencing revealed highest editing rates with pegRNAs mPKU-1.1 (PE2: 19.6%) and mPKU-2.1 (PE2: 19.7%) (Fig. 4a, 4b), but due to lower indel rates of mPKU-2.1 (Fig. 4c, 4d, Extended data fig. 7b) we decided to use this pegRNA for *in vivo* experiments.

**Figure 4:**
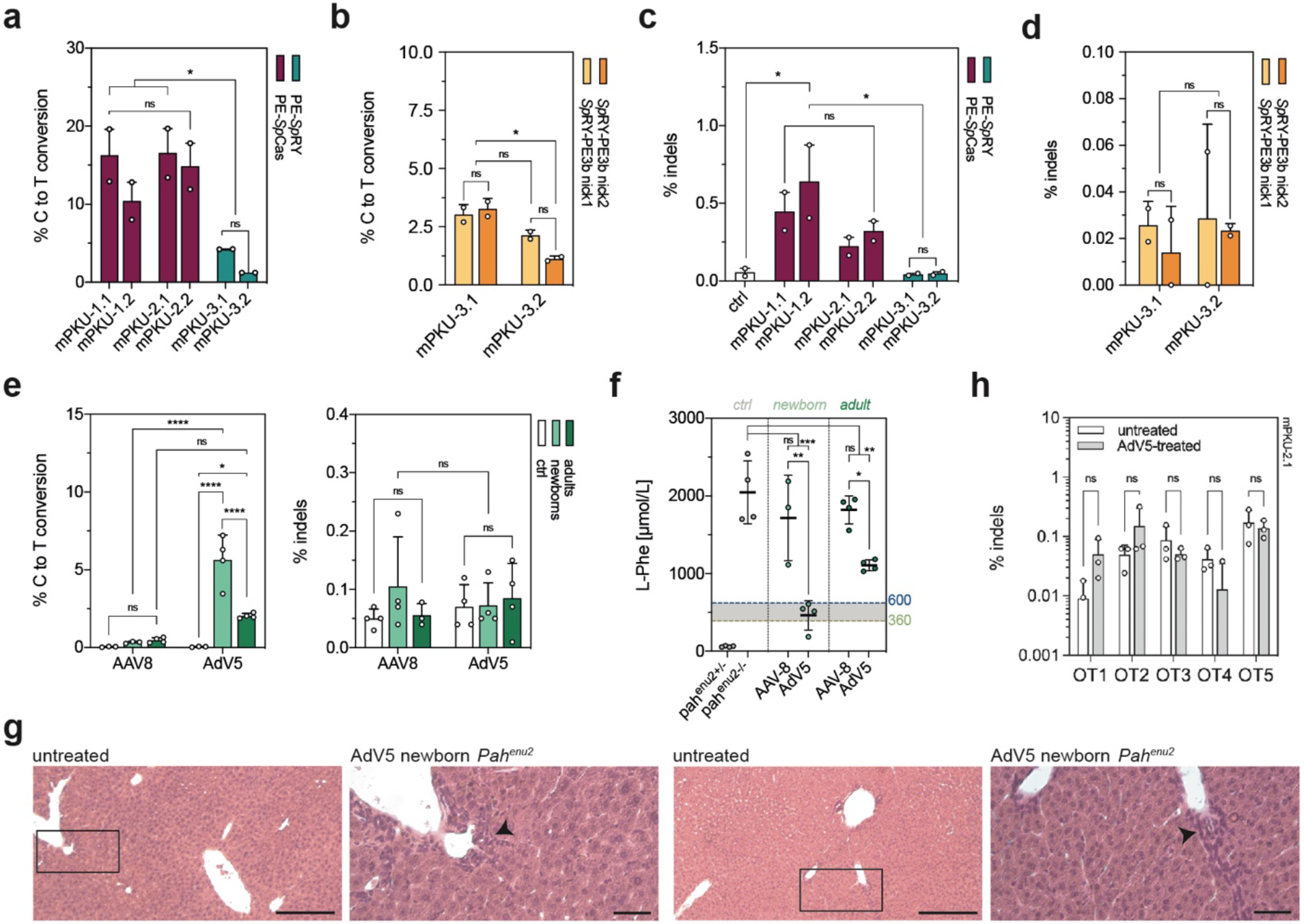
Correction of the *Pah*^*enu2*^ allele *in vivo* in mice using prime editing. (**a-d**) *In vitro* editing rates and indel formation of pegRNAs designed for an *Sp*Cas-(purple) and *Sp*RY-PEs (green) to target the disease-causing *Pah*^*enu2*^ mutation (c.835T>C; p.F263S) on exon 7. Experiments were performed in reporter HEK293T cells in which the mutated exon 7 of the *Pah*^*enu2*^ gene was stably integrated. Data are represented as mean ± range of two independent experiments and were analyzed using a two-way ANOVA with Tukey’s multiple comparisons test (ns, not significant; *\*P>0.05*). (**e**) *In vivo* correction and indel rates in newborn and adult animals after AAV8- or AdV5-mediated delivery of *Sp*Cas-PEs (injected dose of AAV8 in neonates and adults: 1×10^12^ and 2×10^12^ vg; injected dose of AdV5 in neonates and adults: 3×10^10^ and 1.0×10^11^ viral particles, vp). Untreated mice were used as negative controls. Indels are calculated as percentage of sequencing reads with indels around the protospacer region. (**f**) Blood L-Phe levels after *in vivo* prime editing compared to untreated heterozygous and homozygous control animals. L-Phe levels of 360-600 μmol/L are considered therapeutically satisfactory *(54)(55)* (highlighted in grey). (**g**) H&E-stained liver sections of untreated and AdV5-treated mice revealed no differences in mononuclear cell infiltration (arrowhead) and hepatocyte necrosis. Scale bars: 100 μm; 20 μm for higher magnification. (**h**) Deep amplicon sequencing of the top 5 off-target sites for the protospacer of the pegRNA mPKU-2.1, experimentally determined by CHANGE-seq *(39)*, in untreated and AdV5-treated mice (>20’000 reads per site). Data are represented as mean ± s.d. of three animals and were analyzed using a two-way ANOVA with Tukey’s multiple comparisons test (*\*P>0.05; ***P>0.0005; ****P>0.0001*).

### Subhead 6: Restoring physiological L-Phe levels by AdV5-mediated *in vivo* prime editing

To elucidate whether we can reach therapeutic correction of the pathogenic *Pah*^*enu2*^ allele by *in vivo* prime editing, we first generated AAV vectors co-expressing intein-split PE2^Δ*RnH*^ and mPKU-2.1 pegRNA. Systemic delivery in a 1:1 ratio into newborn and adult PKU mice at a dose of 5×10^11^ (newborn) or 1×10^12^ (adult) vg per construct and animal (Extended data fig. 7c), however, led to consistently lower editing rates as compared to the *Dnmt1* locus (< 1%, Fig. 4e). Since AAV transduction rates were comparable in *Dnmt1* and *Pah*^*enu2*^ experiments (Extended data fig. 8a), and both loci were edited with similar efficiencies *in vitro* when stably integrated into the genome of HEK293T cells (Extended data fig. 8b), we speculate that *in vivo* in hepatocytes the *Pah*^*enu2*^ locus is less accessible than the *Dnmt1* locus.

Due to our observations at the *Dnmt1* locus, where AdV5-mediated PE delivery led to an increase in editing rates (Fig. 3b, 3d), we next generated an AdV5 vector expressing unsplit PE2^Δ*RnH*^ together with mPKU-2.1. Systemic i.v. delivery of the vector into adult (tail vein) and neonatal (temporal vein) PKU mice (Extended data fig. 7c) again led to a significant increase in editing compared to AAV-mediated intein-split PE2^Δ*RnH*^ delivery, with correction rates of 2.0 ± 0.2% and 5.6 ±1.6% at the *Pah*^*enu2*^ locus, respectively (Fig. 4e). As for the *Dnmt1* locus also for the *Pah*^*enu2*^ locus the increase in on-target editing rates did not result in indel mutations above untreated control animals (Fig. 4e). Notably, co-expression of a PE3 sgRNA targeting a non-canonical NAG PAM *(37)* 4 bp upstream of the disease-causing mutation led to a trend towards elevated editing rates *in vitro* (19.6 ± 2.6%; Extended data fig. 7a), and in injected newborn animals (10.1 ± 2.7%; Extended data fig. 7d, 7e).

To next assess therapeutic efficacy of AAV8- and AdV5-mediated *in vivo* prime editing of the *Pah*^*enu2*^ locus, we quantified blood L-Phe levels in treated mice. Compared to L-Phe levels of untreated homozygous animals (2046±404 μmol/L), we observed a significant, though not therapeutic *(38)* reduction of L-Phe levels in AdV5-adult mice (1106±70 μmol/L and 1430±468 μmol/L). However, in animals injected with AdV5 as newborns the L-Phe concentration was reduced to therapeutically satisfactory levels of 463±188 μmol/L and to 100±34 μmol/L (Fig. 4f, Extended data fig. 7f), with L-Phe levels maintaining stable over a period of 18 weeks after injection (Extended data fig. 9a). In line with reduced blood L-Phe levels, we also observed restoration of PAH enzyme activity (2.0%-6.0% of wild-type enzyme activity) in liver lysates of treated animals (Extended data figure 9b). Taken together, our data demonstrate that AdV5-mediated *in vivo* prime editing can correct a pathogenic mutation at the *Pah* locus at sufficiently high levels to rescue the PKU phenotype.

### Subhead 7: *In vivo* prime editing does not induce extensive liver damage or off-target mutations

For clinical application of *in vivo* prime editing persistent liver damage and extensive off-target editing triggered by PE expression could be critical limitations. Therefore, we performed histological assessment of liver sections from AdV5- and AAV8-treated animals. In none of the animals we observed an increase in mononuclear immune cell infiltration or hepatocyte necrosis compared to untreated controls (Fig. 4g; Extended data fig. 10a). Likewise, we only detected a very mild elevation of the serum transaminases alanine aminotransferase (ALT) and aspartate aminotransferase (AST) at the timepoints when animals were sacrificed (Extended data fig. 10b). Next, we assessed whether prime editing was restricted to the liver and the targeted *Pah*^*enu2*^ locus. Deep sequencing of the *Pah*^*enu2*^ allele in various tissues of neonatal mice confirmed that substantial prime editing rates were observed the liver (Extended data fig. 10c). In addition, we used CHANGE-seq *(39)* to experimentally identify potential off-target binding sites of the mPKU-2.1 pegRNA (Extended data fig.10d). In line with previous studies that reported higher specificities of PEs compared to CRISPR-Cas9 nucleases *(1, 40)*, targeted amplicon deep sequencing of the top 5 off-target sites for the pegRNA revealed no detectable editing compared to untreated controls (Fig 4h; n=3 mice per group). Likewise, no editing was observed when the top 10 computationally predicted off-target binding sites of mPKU-2.1 were sequenced (Extended data fig. 10e).

Taken together, our data demonstrate that AdV5-mediated prime editing restores the pathogenic *Pah*^*enu2*^ allele *in vivo* without altering liver physiology or inducing off-target editing.

## DISCUSSION

*In vivo* prime editing in somatic tissues holds great potential for therapeutic application in patients with genetic diseases. In our study we demonstrate installation of transition and transversion edits *in vivo* in the liver of mice with high precision. Unlike CRISPR-Cas9 nucleases, PEs work independent of double-strand (ds)DNA break formation. This is an important aspect for safety consideration, as dsDNA breaks can trigger excessive genetic damage, including translocations, inversions, and large deletions *(41–43)*. Another genome editing tool that works independent of HDR are BEs, which have previously been applied to correct disease-causing mutations *in vivo* in somatic tissues in a number of loci *(12–14, 44)*, including the *Pah*^*enu2*^ locus *(13, 16)*. Interestingly, cytidine base editing was more efficient in correcting the PKU-causing point mutation, but unlike prime editing also induced substantial amounts of bystander mutations (up to 61% of edited alleles) and indel mutations (up to 27% of edited alleles) *(16)*. Similar results have also been obtained at other loci^14^, suggesting that excision of the uracil intermediate in combination with DNA nicking frequently leads to dsDNA breaks during the editing process. Thus, PEs might not only be highly valuable for correcting indel mutations and transversion substitutions *(1)*, but also for correcting transition mutations at loci where base editing induces high rates of bystander or indel edits.

In our study, systemic injection of a single dose of AdV5 encoding the prime editing components into neonates resulted in an average of 58% editing at the *Dnmt1* and 8% editing at the *Pah*^*enu2*^ locus. These rates are substantially higher than in recently reported prime editing studies in somatic tissues *(20, 45)*, and led to the restoration of physiological blood-L-Phe levels in *Pah*^*enu2*^ mice. Importantly, correction rates of approximately 10% in hepatocytes should also be sufficient to treat a variety of other genetic liver diseases, including tyrosinemia and urea cycle disorders *(46–48)*. The high vector doses required for efficient editing, however, are a challenge for potential application in patients. Particularly for sites that are less susceptible to prime editing, e.g. due to an unfavorable GC count in the protospacer sequence or reduced locus accessibility, the required AAV or AdV5 concentrations would likely exceed the doses that can be safely applied in humans. Future work should therefore focus on enhancing PE activity in order to sustain clinical viability of *in vivo* prime editing. Examples for such strategies could be to increase nuclear import efficiency of PE through an optimized NLS design *(20)*, or to increase reactivity of the RT domain via directed protein evolution *(49)*.

## MATERIALS AND METHODS

### Generation of plasmids

To generate pegRNA plasmids, annealed spacer, scaffold, and 3’ extension oligos were cloned into pU6-pegRNA-GG-acceptor by Golden Gate assembly as previously described *(1)*. To generate nicking sgRNA plasmids, annealed and phosphorylated oligos were ligated into *BsmBI*-digested lentiGuide-Puro backbone. Sequences of all pegRNAs and nicking sgRNAs are listed in Supplementary Table 1. For the generation of split-intein and orthogonal PEs, inserts were ordered as gBlocks from Integrated DNA Technologies (IDT) and cloned into pCMV-PE2 backbone using HiFi DNA assembly MasterMix (NEB). To generate piggyBac disease reporter plasmids, inserts with homology overhangs for cloning were ordered from IDT and cloned into the pPB-Zeocin backbone using HiFi DNA assembly MasterMix (NEB). To engineer plasmids for virus production, inserts were ordered as gBlocks (IDT) and cloned into AAV backbones using HiFi DNA assembly MasterMix (NEB). All PCR reactions were performed using Q5 High-Fidelity DNA polymerase (New England Biolabs).

pU6-pegRNA-GG-acceptor (Addgene plasmid no. 132777) and pCMV-PE2 (Addgene plasmid no. 132775) were gifts from David Liu. lentiGuide-Puro and PX404 *Campylobacter jejuni* Cas9 were a gift from Feng Zhang (Addgene plasmid no. 52963 and 68338). pCMV-VSV-G was a gift from B. Weinberg (Addgene plasmid no. 8454) and psPAX2 was a gift from D. Trono (Addgene plasmid no. 12260). *Sauri*ABEmax was a gift from Yongming Wang (Addgene plasmid no. 135968).

### Cell culture transfection and genomic DNA preparation

HEK293T (ATCC CRL-3216) and Hepa1-6 (ATCC CRL-1830) cells were maintained in Dulbecco’s Modified Eagle’s Medium (DMEM) plus GlutaMax (Thermo Fisher), supplemented with 10% (v/v) fetal bovine serum (FBS) and 1% penicillin/streptomycin (Thermo Fisher) at 37°C and 5% CO_2_. Cells were maintained at confluency below 90% and seeded on 48-well cell culture plates (Greiner). Cells were transfected at 70% confluency using 1.5 μL of Lipofectamine 2000 (Thermo Fisher) with 375 ng PE, 125 ng pegRNA, and 40 ng sgRNA according to the manufacturer’s instructions. When intein-split PEs were transfected, 375 ng of each PE half was used for transfection. Unless otherwise noted, cells were incubated for 3 days and genomic DNA was isolated by direct lysis.

### Generation of reporter cells by Lentivirus transduction or PiggyBac transposon

For Lentivirus production, HEK293T cells were seeded into T75 flask (Greiner) with Opti-MEM (Thermo Fisher) and transfected at 70% confluency using polyethylenimine (PEI). Briefly, 60 μL PEI (0.1 mg/mL) was mixed with 370 μL Opti-MEM, incubated at room temperature for 5 min, and added to 4.4 μg PAX2, 1.5 μg VSV-G, and 5.9 μg lentiviral vector plasmid (filled up to 430 μL Opti-MEM). Following 20 min incubation at room temperature, cells were transfected. The culture medium was changed one day after transfection. After two days, the cell culture supernatant was harvested and lentiviral particles were purified by filtration (0.20 μm, Sarstedt). Fresh HEK293T cells were subsequently transduced with lentiviral particles in a 24-well cell culture plate (Greiner). Two days after transduction, cells were enriched for 7 days using 2.5 μg/mL Puromycin.

For generation of disease reporter cell lines with the PiggyBac transposon, 30’000 HEK293T cells were seeded into a 24-well culture plate (Greiner) and transfected at 70% confluency using Lipofectamine 2000 (Thermo Fisher) according to the manufacturer’s instructions. Briefly, 1.5 μL Lipofectamine was mixed with 23.5 μL Opti-MEM, incubated at room temperature for 10 min, and added to 225 ng transposon plasmid and 25 ng transposon helper plasmid (filled up to 25 μL Opti-MEM). Following 30 min incubation at RT, cells were transfected. Three days after transfection, cells were enriched for 10 days using 150 μg/mL Zeocin.

### Fluorescence reporter assays and fluorescence-activated cell sorting

Reporter cells were transfected with prime editing tools that are programmed to restore the expression of a fluorescent protein. Cells were incubated for 3 days post-transfection and trypsinized with TrypLE (Gibco). Cells were washed twice with phosphate-buffered saline (PBS) and resuspended in FACS Buffer (PBS supplemented with 2% FBS and 2 mM EDTA). Cell suspensions were filtered through 35 μm nylon mesh cell strainer snap caps (Corning) and kept on ice until analysis. For each sample, 100’000 events were counted on an LSR Fortessa (BD Biosciences) using the FACSDiva software version 8.0.1 (BD Biosciences). Experiments were performed in up to four replicates on different days. Data are reported as mean values ± standard deviation (s.d.). The gating strategy is shown in Extended data Fig. 1.

### AAV and Adenovirus production

All pseudo-typed vectors (AAV2 serotype 8) were produced by the Viral Vector Facility of the Neuroscience Center Zurich. Briefly, AAV vectors were ultracentrifuged and diafiltered. Physical titers (vector genome per mL) were determined using a Qubit 3.0 fluorometer as previously published *(50)*. Identity of the packaged genomes of each AAV vector was confirmed by Sanger sequencing. All human Adenovirus 5 vectors were produced by *ViraQuest, Inc*.

### Animal studies

Animal experiments were performed in accordance with protocols approved by the Kantonales Veterinäramt Zürich and in compliance with all relevant ethical regulations. *Pah*^*enu2*^ and C57BL/6J mice were housed in a pathogen-free animal facility at the Institute of Pharmacology and Toxicology of the University of Zurich. Mice were kept in a temperature- and humidity-controlled room on a 12-h light-dark cycle. Mice were fed a standard laboratory chow (Kliba Nafag no. 3437 with 18.5% crude protein) and genotyped at weaning. Heterozygous *Pah*^*enu2*^ littermates were used as controls for physiological L-Phe levels in the blood (<120 μM). For sampling of blood for L-Phe determination, mice were fasted for 3-4 h and blood was collected from the tail vein. Unless otherwise noted, newborn animals (P1) received 3.0×10^10^ (Adenovirus, 30 μL total) or 5×10^11^ (AAV, 30 μL total) vg per animal and construct via temporal vein. Adult mice were injected with 1.0×10^11^ (Adenovirus, 150 μL total) per animal or 1×10^12^ (AAV, 150 μL total) vg per animal and construct via tail vein. Average weight of neonatal and adult mice (5 weeks) was 1.5 g and 20 g, respectively. Newborn mice were euthanized 4 weeks after injection. Adult mice were euthanized 4 weeks (Adenovirus) or 12 weeks (AAV) after injection if not stated otherwise.

### Primary hepatocyte isolation

Primary hepatocytes were isolated using a two-step perfusion method. Briefly, pre-perfusion with HANKS’s buffer (HBSS supplemented with 0.5 mM EDTA, 25 mM HEPES) was performed by inserting the cannula through the superior vena cava and cutting the portal vein. Next, livers were perfused at low flow for approximately 10 min with Digestion Buffer (low glucose DMEM supplemented with 1 mM HEPES) containing freshly added Liberase TM (32 μg/mL; Roche). Digestion was stopped using Isolation Buffer (low glucose DMEM supplemented with 10% FBS) and cells were separated from the matrix by gently pushing with a cell scraper. The cell suspension was filtered through a 100 μm filter (Corning) and hepatocytes were purified by two low speed centrifugation steps (50×g for 2 min).

### RNA isolation and RT-qPCR

RNA was isolated from shock frozen liver samples using the RNeasy Mini kit (Qiagen) according to the manufacturer’s instructions. RNA was reverse transcribed to cDNA using random primers and GoScript™ Reverse Transcriptase kit (Promega). RT-qPCR was performed using Firepol qPCR Master Mix (Solis BioDyne) and analyzed by 7900HT Fast Real-Time PCR System (Applied Biosystems). Fold changes were calculated using the delta Ct method. Used primers are listed in Suppl. Table 4.

### Western Blot

Proteins were isolated from transfected HEK293T cells or transduced primary hepatocytes of treated and untreated animals. Briefly, cells were lysed in RIPA buffer, supplemented with protease inhibitors (Sigma-Aldrich). Protein amounts were determined using the Pierce™ BCA Protein Assay Kit (Thermo Fisher). Equal amounts of protein (*in vitro* samples: 3 μg; *in vivo* samples: 80 μg) were separated by SDS-PAGE (Thermo Fisher) and transferred to a 0.45 μm nitrocellulose membrane (Amersham). Membranes were incubated with mouse anti-Cas9 (1:1’000; Cat. No. #14697T; Cell Signaling) and rabbit anti-GAPDH (1:10’000; Cat. No. ab181602; abcam). Signals were detected by fluorescence using IRDye-conjugated secondary antibodies (Licor).

### Amplification for deep sequencing

Genomic DNA from mouse liver tissues were isolated from whole liver lysate by direct lysis. Locus-specific primers were used to generate targeted amplicons for deep sequencing. First, input genomic DNA was amplified in a 20 μL reaction for 25 cycles using NEBNext High-Fidelity 2× PCR Master Mix (NEB). PCR products were purified using AMPure XP beads (Beckman Coulter) and subsequently amplified for 8 cycles using primers with sequencing adapters. Approximately equal amounts of PCR products from each sample were pooled, gel-purified and quantified using a Qubit 3.0 fluorometer and the dsDNA HS assay kit (Thermo Fisher). Paired-end sequencing of purified libraries was performed on an Illumina Miseq.

### HTS data analysis

Sequencing reads were demultiplexed using MiSeq Reporter (Illumina). Amplicon sequences were aligned to their reference sequences using CRISPResso2 *(51)*. Prime editing efficiencies were calculated as percentage of (number of reads containing only the desired edit)/(number of total reads). Indel yields were calculated as percentage of (number of indel-containing reads)/(total reads).

### Guide-dependent off-target prediction and analysis using CHANGE-seq

For CHANGE-seq, the protospacer of the pegRNA was first tested for functionality: a 599 bp long construct was PCR amplified from genomic DNA isolated from the tail of a *Pah*^*enu2*^ homozygous animal using GoTaq G2 HotStart Green Master Mix (Promega). Primers are listed in Suppl. Table 2. The library was prepared as previously described *(39)*. Data was processed using v1.1 of the CIRCLE-Seq analysis pipeline (https://github.com/tsailabSJ/circleseq) with parameters: “window_size: 3; mapq_threshold: 50; start_threshold: 1; gap_threshold: 3; mismatch_threshold: 15; merged_analysis: True; variant_analysis: True”. The top 5 off-target sites for the protospacer of the pegRNA were selected for targeted amplicon deep sequencing and covered by at least 20’000 reads per site.

### PAH enzyme activity assay

Whole liver extracts were analyzed using isotope-dilution liquid chromatography-electrospray ionization tandem mass spectrometry (LC-ESI-MS/MS) according to a previously published method *(52)*.

### Quantification of Phenylalanine in blood

Amino acids were extracted from a 3.2 mm dried blood sample using the Neomass AAAC Plus newborn screening kit (Labsystems Diagnostics, Helsinki, Finland). A UHPLC Nexera X2 coupled to an LCMS-8060 Triple Quadrupole mass spectrometer with electrospray ionization (Shimdazu, Kyoto, Japan) was used for the quantitative analysis of Phenylalanine. Labsolutions and Neonatal Solutions software (Shimdazu, Kyoto, Japan) were used for data acquisition and data analysis.

### Partial (70%) hepatectomy

0.1 mg/kg bodyweight of buprenorphine was administered into the mice subcutaneously 30 min before starting the operative procedure. Animals were anesthetized with isoflurane: 5% isoflurane with 1000 mL/min in 100% O_2_. During the surgical procedure, anesthesia was maintained at 1.5-2.5% isoflurane with 400 mL/min in 100% O_2_. Correct level of anesthesia was verified by toe-pinch reflex. The animals were placed on a warming surface for maintenance of body temperature. The incision site was disinfected with antiseptic, povidone-iodine solution (Braunol, Braun). A midline abdominal skin and muscle incision (about 3 cm long) was performed to expose the abdominal cavity. The liver was freed from ligaments. After cholecystectomy (Prolene, 8/0), the left lateral lobe was ligated with a 6/0 silk thread (Fine Science Tools) and resected. Successful ligation was confirmed by observing a color change of the lobe. The median lobe was ligated in two steps (silk 6/0) and resected. For closing the peritoneum and the abdomen, VICRYL® 5-0 Suture (Ethicon) was used.

### Histology

Livers were fixed using 4% paraformaldehyde (PFA, Sigma-Aldrich), followed by ethanol dehydration and paraffinization. Paraffin blocks were cut into 5 μm thick sections, deparaffinized with xylene, and rehydrated. Sections were HE-stained and examined for histopathological changes.

### Immunofluorescence

Antigen retrieval was performed on deparaffinized and rehydrated 5 μm thick liver sections using HistosPro Microwave histoprocessor. Slides were blocked using PBS supplemented with 2% Normal-donkey serum (Abcam, ab7475) and 0.1% Triton-X for 30 min. Slides were incubated with primary antibody at 4°C overnight using rabbit-anti-CK19 (Abcam, ab52625, dilution 1:500) and mouse-anti-Ki67 (Agilent, M7248, dilution 1:100). Goat-anti-mouse-488 and Goat-anti-rabbit-568 were used as secondary antibodies and sections were counterstained with DAPI. Mounting was performed using Fluorescence Mounting Medium (Agilent, S302380-2). Confocal Images were taken with a Zeiss LSM 700. Ki67-positive cells per area were quantified using the cell counter plugin of FIJI *(53)*.

### Statistical analysis

*A priori* power calculations to determine sample sizes for animal experiments were performed using the R ‘pwr’ package. Statistical analyses were performed using GraphPad Prism 9.0.0. for MacOS. Data are represented as biological replicates and are depicted as mean ± s.d. as indicated in the corresponding figure legends. Likewise, sample sizes and the statistical tests used are described in detail in the respective figure legends. For all analyses, *P<0.05* values were considered statistically significant.

## Supplementary Materials

### Supplementary figures

Fig. S1. Gating strategies for HEK293T reporter cell lines.

Fig. S2. Optimization of intein-split PEs for dual AAV-mediated delivery.

Fig. S3. Orthogonal PE systems display low editing efficiencies in HEK293T cells. Fig. S4. *In vitro* and *in vivo Dnmt1* editing rates and indel formation over time.

Fig. S5. Effects of cell proliferation on *Dnmt1* editing rates. Fig. S6. *In vitro* and *in vivo* reconstitution of intein-split PEs.

Fig. S7. *In vitro* and *in vivo* correction of the *Pah*^*enu2*^ locus using *Sp*Cas-PEs.

Fig. S8. Comparative analysis of AAV transduction efficiency and prime editing reagents for *Pah*^*enu2*^ and *Dnmt1* target sites.

Fig. S9. AdV5-mediated prime editing results in stable editing at the *Pah*^*enu2*^ target site and restoration of PAH activity.

Fig. S10. *In vivo* prime editing does not induce extensive liver damage or off-target editing.

### Supplementary tables

Table S1. Oligos used for deep sequencing.

Table S2. Oligos used for cloning of pegRNAs and nicking gRNAs.

Table S3. Nucleotide sequences of HTS amplicons.

Table S4. Oligos used for RT-qPCR.

Table S5. Amino acid sequences of PE constructs.

### Supplementary Data

SD. 1. Complete images of Western Blots.

## Acknowledgments

The Functional Genomics Center Zurich and ZMB are acknowledged for technical support and access to instruments at the University of Zurich and ETH Zürich. Members of the Schwank lab are acknowledged for discussions.

## Author contributions

Conceptualization: DB, TR, LV, GS

Investigation: DB, TR, LV, LS, NM, EI, NR, HMGC

Visualization: DB, TR, LV,

Funding acquisition: DB, BT, GS

Supervision: GS

Writing – original draft: DB, GS

Writing – review & editing: DB, TR, LV, GS

## Data and materials availability

No custom code has been used in this study. Illumina sequencing data will be submitted to Gene Expression Omnibus (GEO) database and will be made publicly accessible prior to publication. The authors declare that all other data supporting the findings of this study are available within the paper and its extended data files or upon reasonable request.

