## Supplementary Files for "Treatment of a metabolic liver disease by *in vivo* prime editing in mice"

**Affiliations:**

McGovern Institute for Brain Research at MIT, 46-5023C, 43 Vassar St. Cambridge, Middlesex County 02139

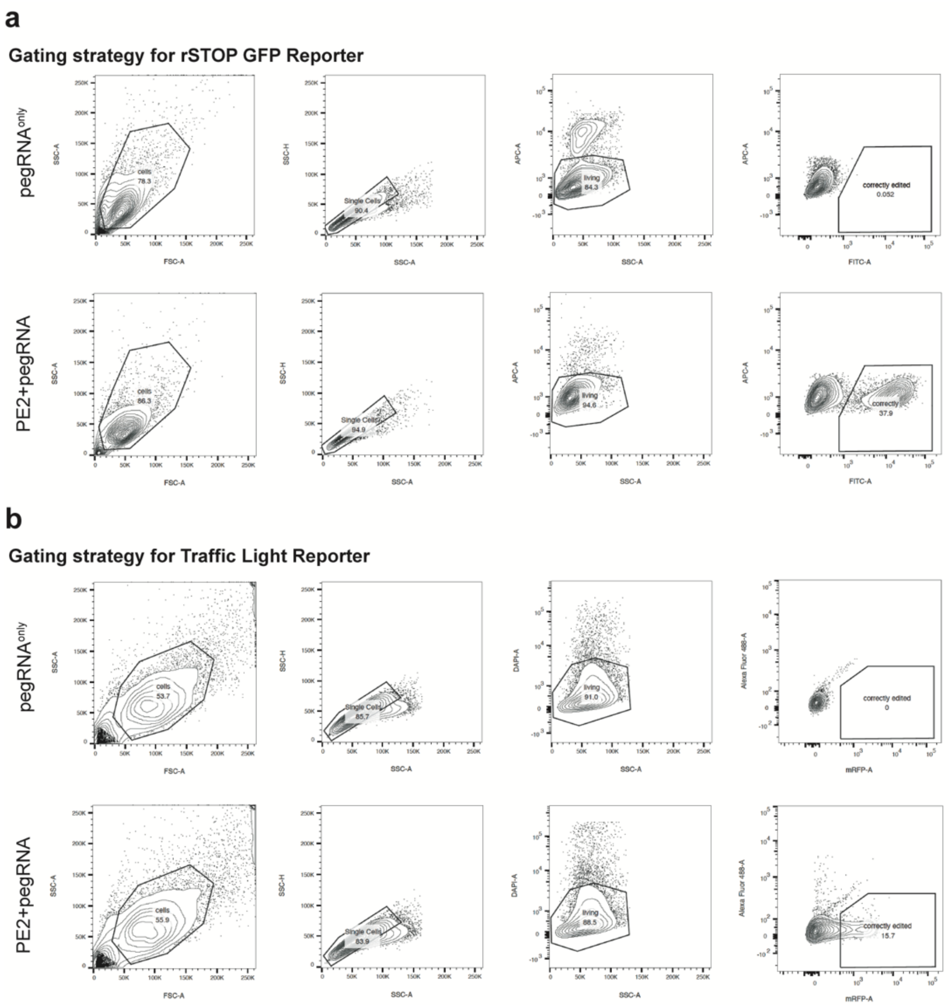


**Figure S1: Gating strategies for HEK293T reporter cell lines.** (**a, b**) Fluorescent signals are scored as successful editing events.

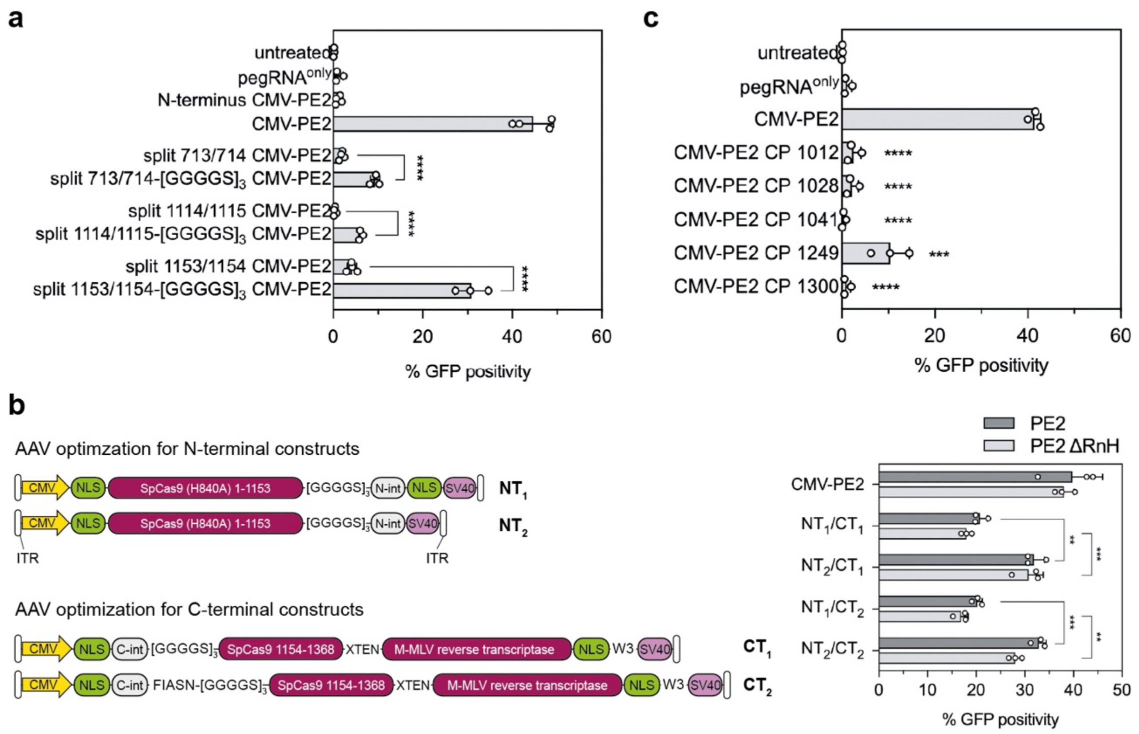


**Figure S2: Optimization of intein-split PEs for dual AAV-mediated delivery.** (**a**) The effect of a Glycine-Serine ([GGGGS]3)-linker on prime editing efficiencies at different split sites. Statistical analysis was performed using a two-tailed student's t-test (*****P>0.0001*). (**b**) Schematic maps of N- and C-terminal intein-split PE halves for AAV delivery. Maps are not to scale. Positioning of NLSs alters the performance of intein-split PE2-p.1153. Data were analyzed using a two-way ANOVA with Tukey's multiple comparisons test (***P>0.005*; ****P>0.0005*). (**c**) Editing efficiencies of different circular permutant (CP) PE2 compared to full-length PE2. Amino acid positions, at which N- and C-termini were interchanged, are indicated. Data were analyzed using a two-tailed student's t-test (****P>0.0005*; *****P>0.0001*). Data in (a-c) are represented as mean ± s.d. of at least three independent biological replicates. ITR, inverted terminal repeat sequences; W3, Woodchuck Hepatitis Virus post-transcriptional regulatory element; SV40, polyadenylation signal.

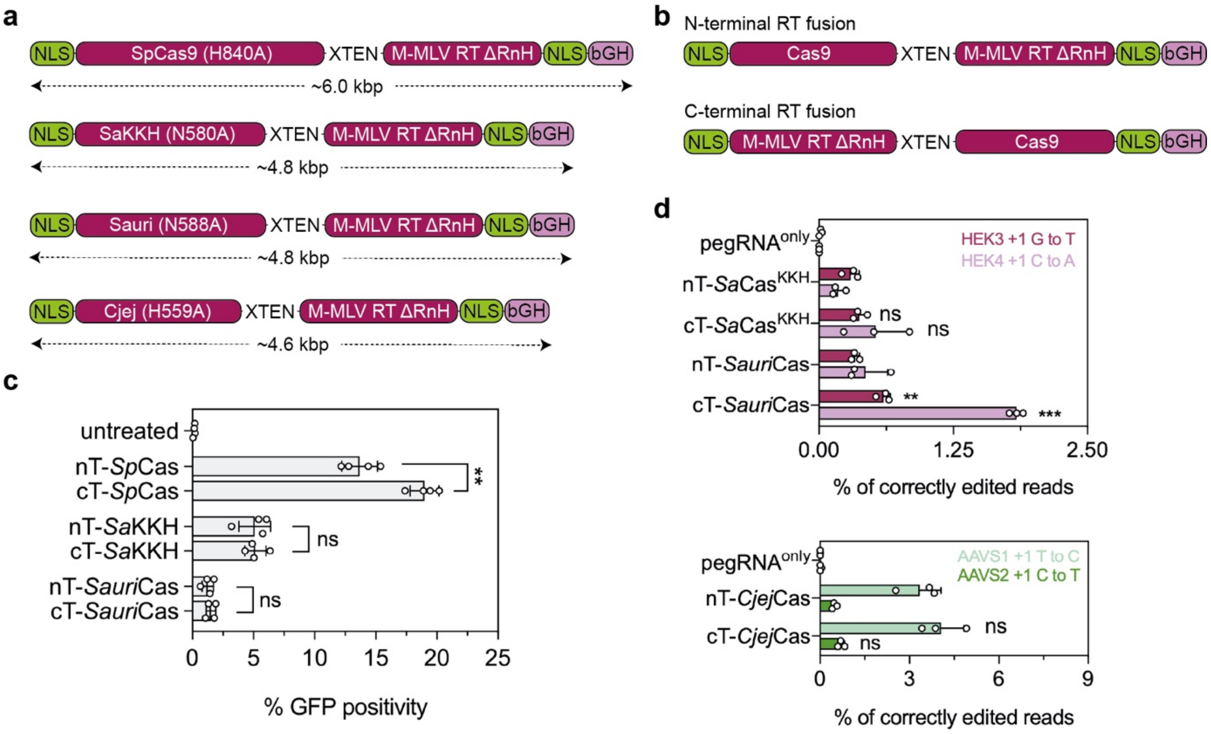


**Figure S3: Orthogonal PE systems display low editing efficiencies in HEK293T cells.** (**a, b**) Schematic maps and sizes of orthogonal PE2$^{\Delta RnH}$ where the RT is fused to Cas9 from *S. aureus*, *S. auricularis*, and *C. jejuni*. (**c, d**) Editing efficiencies of the different PE2$^{\Delta RnH}$ variants in rSTOP reporter cells (c) and on endogenous sites (d). One pegRNA was designed (PBS: 12-14 nucleotides, RT: 15-20 nucleotides) and tested for each locus. Data are represented as mean ± s.d. of at least three independent biological replicates. Data were analyzed using a two-tailed student's t-test (ns, not significant; ***P<0.005; ***P<0.001*). nT, N-terminal RT fusion; cT, C-terminal RT fusion.

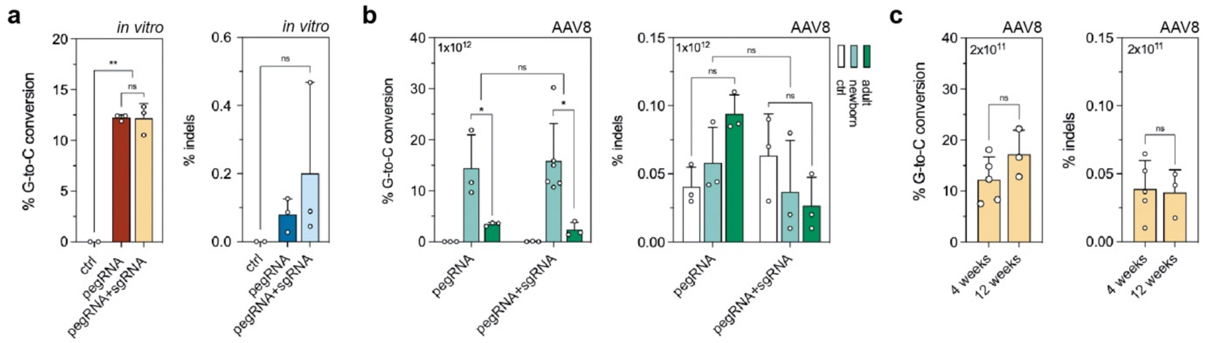


**Figure S4: *In vitro* and *in vivo Dnmt1* editing rates and indel formation over time.** (**a**) G-to-C transversion rates and indel formation for the pegRNA and the pegRNA+sgRNA at the endogenous *Dnmt1* locus in Hepa1-6 cells. (**b**) *In vivo* editing rates and indel formation in neonates and adults treated with pegRNA or pegRNA+sgRNA, respectively (injected AAV dose: $1x10^{12}$). (**c**) *In vivo* editing rates and indel formation in neonates at 4 weeks and 12 weeks. Data are represented as mean ± s.d. (n= 3-6 replicates/mice per group) and were analyzed using two-way ANOVA with Tukey's multiple comparisons test (ns, not significant; ***P>0.005*).

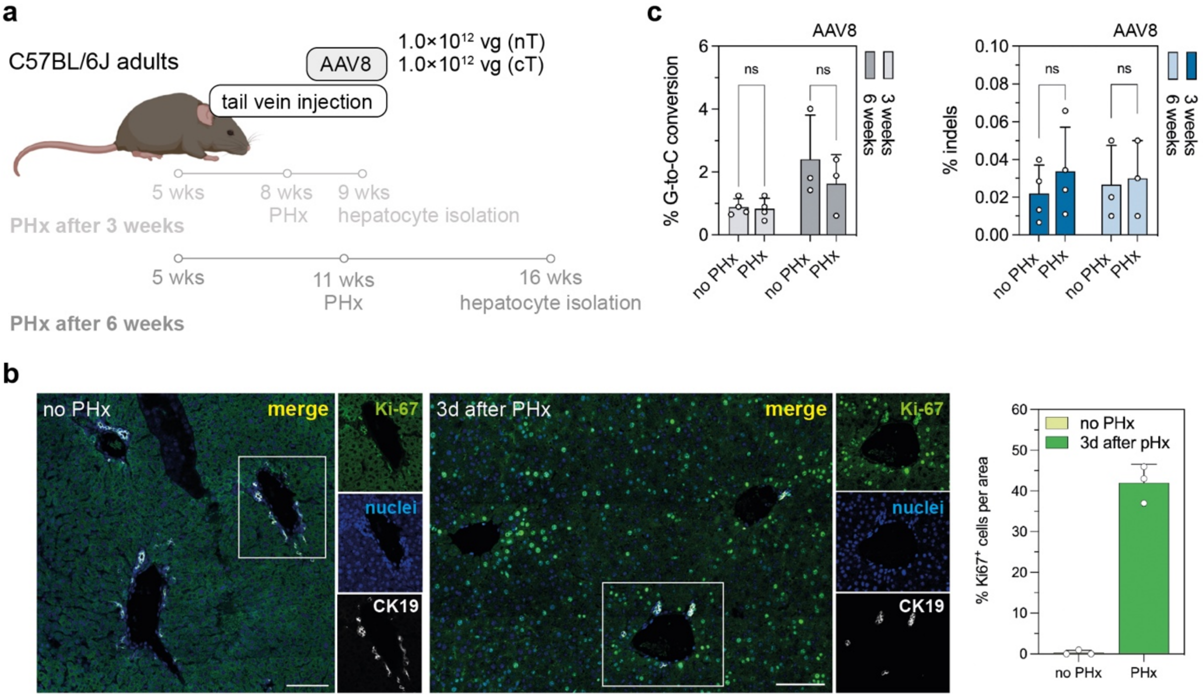


**Figure S5: Effects of cell proliferation on *Dnmt1* editing rates.** (**a**) Schematic outline of the experimental setup for AAV-mediated delivery and partial hepatectomy (PHx) in adult mice (wks, weeks). (**b**) Ki-67 staining of liver cryo-sections from control (no PHx) and PHx mice (3d after surgery). Three areas were randomly selected across the liver for quantification of proliferating cells per field of view (n=3). (**c**) Effect of PHx, performed at 3 or 6 weeks, on on-target prime editing rates and indel levels in adult mouse livers. Data are represented as mean ± s.d. (n= 3-4 replicates/mice per group) and were analyzed using two-way ANOVA with Tukey's multiple comparisons test (ns, not significant; $^{**}P>0.005$).

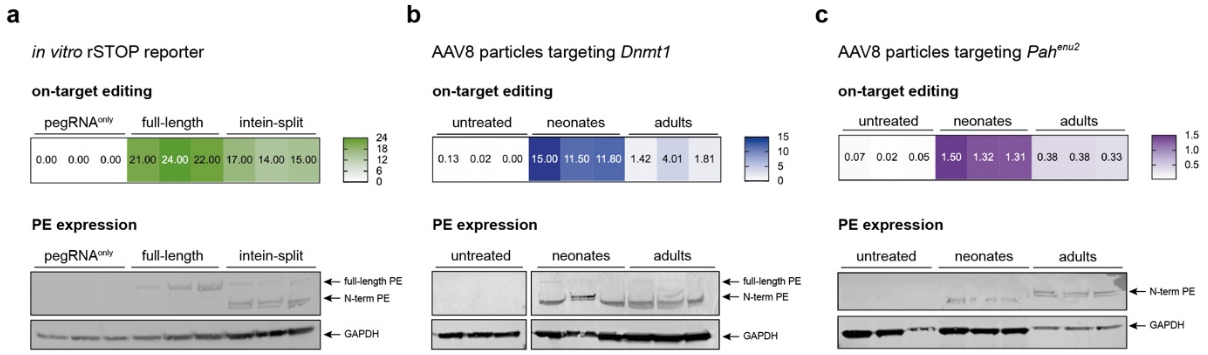


**Figure S6: *In vitro* and *in vivo* reconstitution of intein-split PEs.** (**a**) *In vitro* reconstitution of the intein-split PE and on-target editing in the rSTOP reporter. (**b**) *In vivo* reconstitution of intein-split PE2, delivered by AAV8, in neonatal and adult mice. Prime editor expression was analyzed using a monoclonal antibody targeting the N-terminal part of *Sp*Cas9 (Cell Signaling). Data are represented as mean (n=3 mice per group).

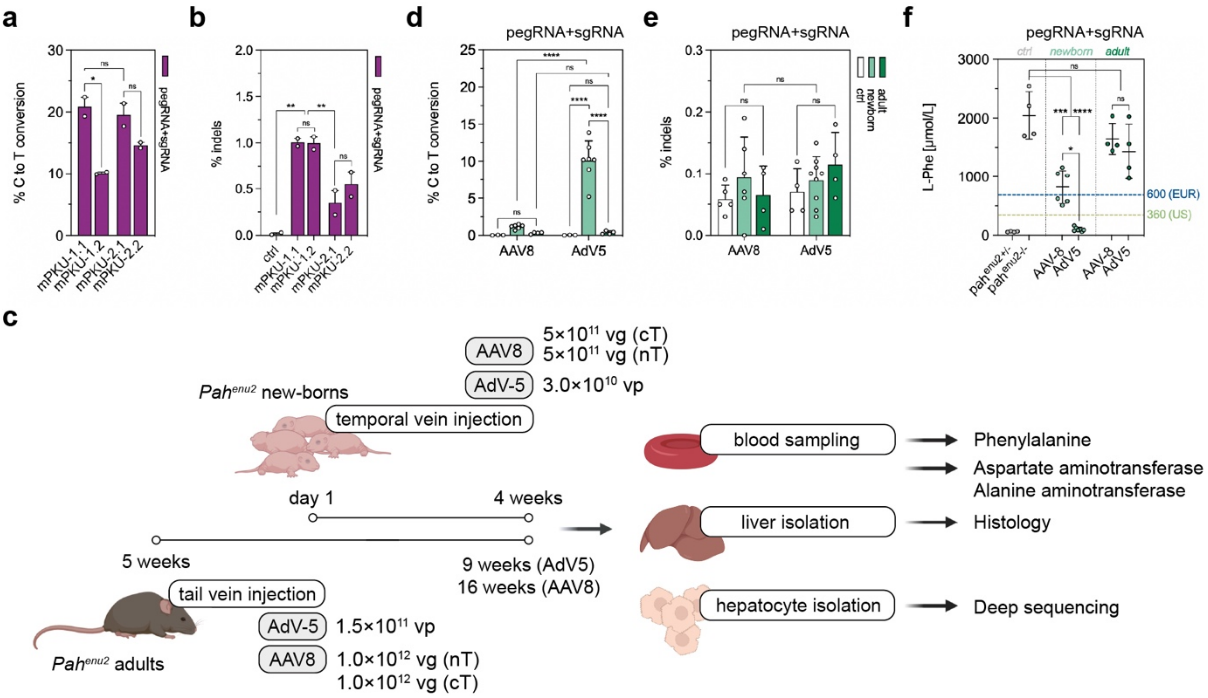


**Figure S7: *In vitro* and *in vivo* correction of the *Pah*$^{enu2}$ locus using *Sp*Cas-PEs.** (**a**, **b**) *In vitro* correction (a) and indel rates (b) of different pegRNA protospacers combined with an additional sgRNA. Percentage of sequencing reads with indels around the protospacer regions were determined by deep amplicon sequencing. pegRNA plasmids were transfected as negative controls. Experiments were performed in reporter HEK293T cells, in which the mutated exon 7 of the *Pah*$^{enu2}$ gene was stably integrated. Data are represented as mean ± range from 2 independent experiments and were analyzed using two-way ANOVA with Tukey's multiple comparisons test (ns, not significant). (**c**) Schematic outline of the experimental setup for AAV8- and AdV5-mediated treatment in newborn and adult PKU mice. (**d, e**) *In vivo* correction (d) and indel rates (e) in newborn and adult animals after AAV8- and AdV5-mediated delivery. Untreated mice were used as negative controls. Percentage of sequencing reads supporting indel formation at the target locus in untreated, AAV8-, and AdV5-treated animals were determined by targeted amplicon sequencing. (**f**) Blood L-Phe levels after *in vivo* prime editing compared to untreated, heterozygous, and homozygous control animals. L-Phe levels of 360-600 µmol/L are considered therapeutic (highlighted in grey). Data are represented as mean ± s.d. (n=3-6 mice per group) and were analyzed using a two-way ANOVA with Tukey's multiple comparisons test (ns, not significant; **P<0.05; ** P<0.005; ***P<0.0005; ****P<0.0001*).

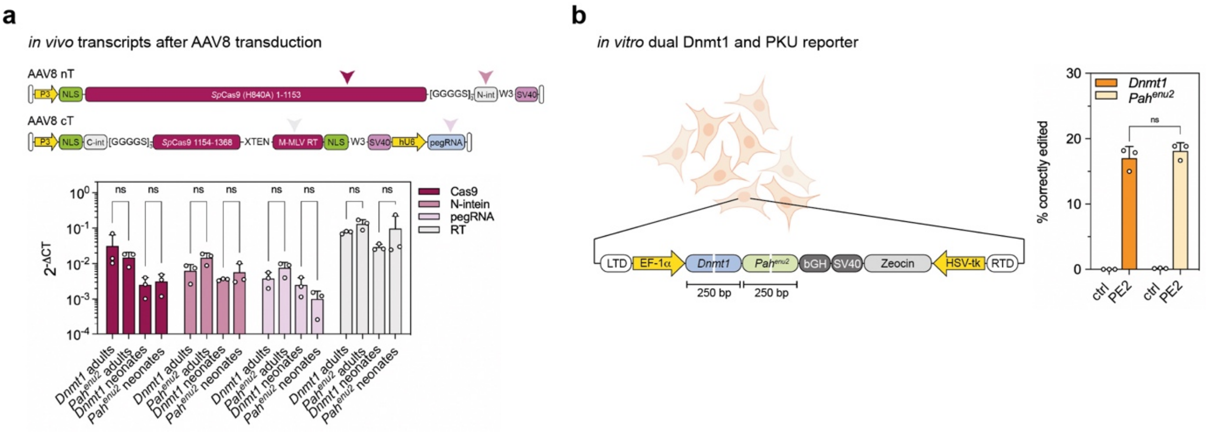


**Figure S8: Comparative analysis of AAV transduction efficiency and prime editing reagents for *Pah*$^{enu2}$ and *Dnmt1* target sites.** (**a**) Transcript analyses of intein-split PE2, delivered by AAV8, in neonatal and adult mice. Primer binding sites on N- (nT) and C-terminal (cT) AAV8 constructs are indicated by arrowheads. Transcript levels were normalized to the housekeeping gene Rplp0. Data are represented as mean ± s.d. (n=3 mice per group) and were analyzed using a two-way ANOVA with Tukey's multiple comparisons test (ns, not significant). (**b**) Comparison of *in vitro* editing rates of the pegRNAs, that were used for *in vivo* experiments, using a dual HEK293T Dnmt1 and PKU reporter cell line, generated using the PiggyBac transposons. pegRNA-binding sites are indicated as white lines. Prime editing efficiencies at both target sites quantified by deep amplicon sequencing. Data are represented as mean ± s.d. of three independent replicates and were analyzed using a two-way ANOVA with Tukey's multiple comparisons test (ns, not significant; $**P>0.005$).

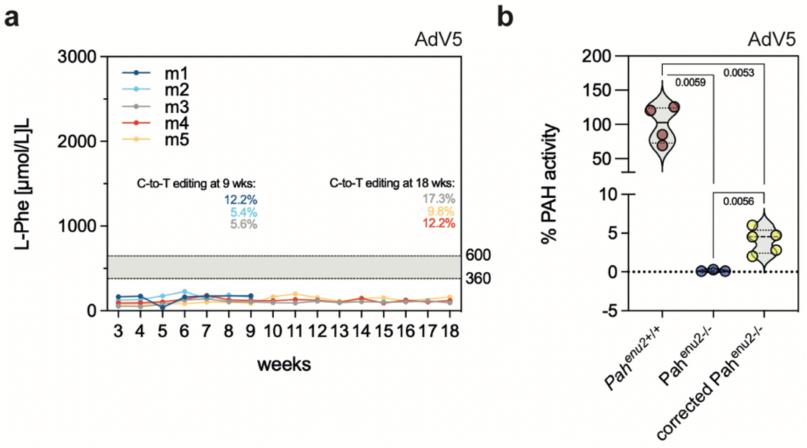


**Figure S9: AdV5-mediated prime editing results in stable editing at the *Pah$^{enu2}$* target site and restoration of PAH activity.** (**a**) Blood L-Phe levels in neonatal mice over 9 weeks. Editing level of the corresponding mice are color-coded and indicated in the top right box. (**b**) Enzymatic activity of PAH at experimental endpoints (4 weeks). Data are represented as mean ± s.d. (n=3-7 mice per group) and were analyzed using a two-way ANOVA with Tukey's multiple comparisons test (ns, not significant; exact *P*-values are indicated in panel c).

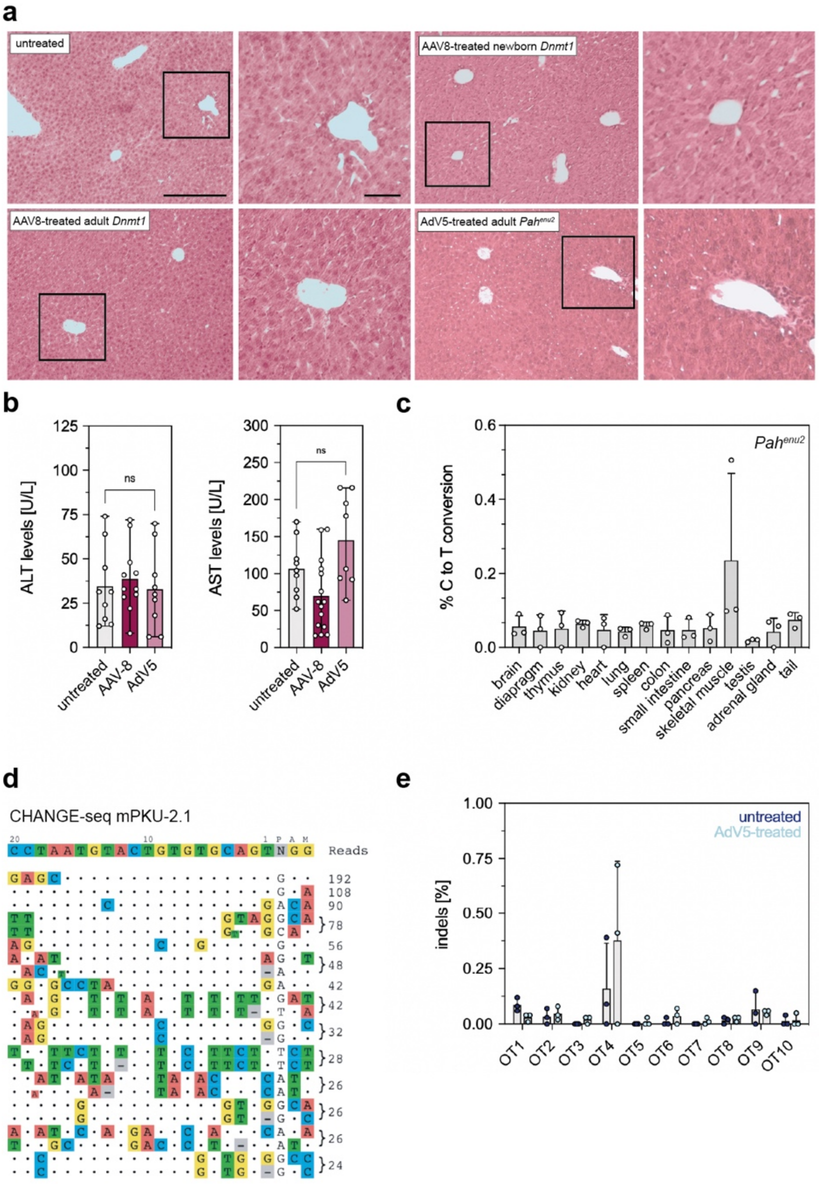


**Figure S10: *In vivo* prime editing does not induce extensive liver damage or off-target editing.** (**a**) Representative H&E-stained liver sections of untreated, AAV8-, and AdV5-treated mice. Vectors were injected into neonatal and adult mice and tissues were analyzed after 4 (AdV5) and 12 weeks (AAV8), respectively. Scale bars, 100 µm (overview) and 50 µm (magnified view). (**b**) Serum transaminases alanine aminotransferase (ALT) and aspartate aminotransferase (AST) at experimental end points of untreated, AAV8-, and AdV5-treated mice (n=8-15 mice per group). (**c**) Deep sequencing of the *Pah*$^{enu2}$ locus in off-target tissues of AdV5-treated neonates. (**d**) Off-targets for the protospacer of the pegRNA mPKU-2.1 were experimentally identified by CHANGE-seq. (**e**) Deep sequencing of 10 computationally predicted off-targets in untreated and AdV5-treated mice (n=3 mice per group). Data are represented as mean ± s.d. (n=3-6 mice per group) and were analyzed using a two-way ANOVA with Tukey's multiple comparisons test (ns, not significant).

**Table 1: Oligos used for deep sequencing.**

| oligo name | oligo sequence |
|---|---|
| HTS_adrb1-fwd | CTTTCCCTACACGACGCTCTTCCGATCTNNNNNNCCAGCATTGAGACCCTGTGT |
| HTS_adrb1-rev | GGAGTTCAGACGTGTGCTCTTCCGATCTNNNNNNCATGAGGATGGGCAGGAAGG |
| HTS_app-fwd | CTTTCCCTACACGACGCTCTTCCGATCTNNNNNNTGGGTAGGCTTTGTCTTACA |
| HTS_app-rev | GGAGTTCAGACGTGTGCTCTTCCGATCTNNNNNNATATCCTGAGTCATGTCGGA |
| HTS_eIF2B4_1-fwd | CTTTCCCTACACGACGCTCTTCCGATCTNNNNNNGTGCTGGAATTCCTAAAAGG |
| HTS_eIF2B4_1-rev | GGAGTTCAGACGTGTGCTCTTCCGATCTNNNNNNGTGATCACCAGATCCACGAG |
| HTS_eIF2B4_2-fwd | CTTTCCCTACACGACGCTCTTCCGATCTNNNNNNGGATCCACTAGTAACGGCCG |
| HTS_eIF2B4_2-rev | GGAGTTCAGACGTGTGCTCTTCCGATCTNNNNNNTCCCTGGAGAGTTCCTCACT |
| HTS_otc-fwd | CTTTCCCTACACGACGCTCTTCCGATCTNNNNNNGGGAGGACACCCTTCCTTTC |
| HTS_otc-rev | GGAGTTCAGACGTGTGCTCTTCCGATCTNNNNNNCAGTCCCTACCTGTGCCCAC |
| HTS_gabaR1α-fwd | CTTTCCCTACACGACGCTCTTCCGATCTNNNNNNCGTCAAAGTTGGAAGGATGA |
| HTS_gabaR1α-rev | GGAGTTCAGACGTGTGCTCTTCCGATCTNNNNNNTATTCATAGGACCGCCATCC |
| HTS_Dntm1-fwd | CTTTCCCTACACGACGCTCTTCCGATCTNNNNNNGTCTTCCCCCACTCTCTTGC |
| HTS_Dntm1-rev | GGAGTTCAGACGTGTGCTCTTCCGATCTNNNNNNCCCCCAATATATGCCTCGGC |
| HTS_PKU-fwd | CTTTCCCTACACGACGCTCTTCCGATCTNNNNNNCCGTCCTGTTGCTGGCTTAC |
| HTS_PKU-rev | GGAGTTCAGACGTGTGCTCTTCCGATCTNNNNNNTGAGCATCCATTGTGGTTGG |
| HTS_OT1peg-PKU-fwd | CTTTCCCTACACGACGCTCTTCCGATCTGGTTTCCCATTTCCCATCATCATT |
| HTS_OT1peg-PKU-rev | GGAGTTCAGACGTGTGCTCTTCCGATCTCTGGGAAGTGTGTACATGTATGG |
| HTS_OT2peg-PKU-fwd | CTTTCCCTACACGACGCTCTTCCGATCTAGCATGTATGGGTTCCGAGG |
| HTS_OT2peg-PKU-rev | GGAGTTCAGACGTGTGCTCTTCCGATCTGAAACCAGTTTCAGCACGGTC |
| HTS_OT3peg-PKU-fwd | CTTTCCCTACACGACGCTCTTCCGATCTATTAGGGAGGAGGGTAGAAGTGT |
| HTS_OT3peg-PKU-rev | GGAGTTCAGACGTGTGCTCTTCCGATCTGCAGTACATGAGCTTCCGC |
| HTS_OT4peg-PKU-fwd | CTTTCCCTACACGACGCTCTTCCGATCTCAGGGCAAGCAGGTAGATGT |
| HTS_OT4peg-PKU-rev | GGAGTTCAGACGTGTGCTCTTCCGATCTACCTGTCGCCCTTCAGTTTG |
| HTS_OT5peg-PKU-fwd | CTTTCCCTACACGACGCTCTTCCGATCTAGTATACCACTGTTTGTGCATTCC |
| HTS_OT5peg-PKU-rev | GGAGTTCAGACGTGTGCTCTTCCGATCTGCAAGTACGCTGCACACAAT |

**Table 2: Oligos used for cloning of pegRNAs and nicking gRNAs.**

| oligo name | oligo sequence |
|---|---|
| rSTOP-a_spacer-fwd | CACCGAgccaccatgGGACTAGAGTGTTTT |
| rSTOP-a_spacer-rev | CTCTAAAACACTCTAGTCCcatggtggcTC |
| rSTOP-a_RT/PBS-fwd | GTGCGGGTACAATCCCACTCTAGTCCCATGGTGG |
| rSTOP-a_RT/PBS-rev | AAAACCACCATGGGACTAGAGTGGGATTGTACCC |
| rSTOP-b_spacer-fwd | CACCGcgtgagTCTAGAgccaccatGTTTT |
| rSTOP-b_spacer-rev | CTCTAAAACatggtggcTCTAGActcacgC |
| rSTOP-b_RT/PBS-fwd | GTGCATACTGAGGGGTACAATCCCACTCTAGTCCCATGGTGGCTCTAGACT |
| rSTOP-b_RT/PBS-rev | AAAAAGTCTAGAGCCACCATGGGACTAGAGTGGGATTGTACCCCTCAGTAT |
| TLR-a_spacer-fwd | CACCgggcgaggagctgttcaccgGTTTT |
| TLR-a_spacer-rev | CTCTAAAACcggtgaacagctcctcgccc |
| TLR-a_RT/PBS-fwd | GTGCgggcaccacccgtgaacagctcctc |
| TLR-a_RT/PBS-rev | AAAAgaggagctgttcacgggtggtgccc |
| TLR-b_spacer-fwd | CACCgtagcccagggtggtcacgaGTTTT |
| TLR-b_spacer-rev | CTCTAAAACtcgtgaccaccctgggctac |
| TLR-b_RT/PBS-fwd | GTGCgccctggcccacccgtgaccaccctggg |
| TLR-b_RT/PBS-rev | AAAAcccagggtggtcacgggtgggccagggc |
| adrb1_spacer-fwd | CACCGAGTGTGGGCCATCTCGGCGTGTTTT |
| adrb1_spacer-rev | CTCTAAAACACGCCGAGATGGCCCACACTC |
| adrb1_RT/PBS-fwd | GTGCGGACACCAACACCGAGATGGC |
| adrb1_RT/PBS-rev | AAAAGCCATCTCGGTGTTGGTGTCC |
| app_spacer-fwd | CACCGGAGATCTCTGAAGTGAAGAGTTTT |
| app_spacer-rev | CTCTAAAACTCTTCACTTCAGAGATCTCC |
| app_RT/PBS-fwd | GTGCATTCTGTATCCATCTTCACTTCAGA |
| app_RT/PBS-rev | AAAATCTGAAGTGAAGATGGATACAGAAT |
| eIF2B4-1_spacer-fwd | CACCGAGACCAGATTCAACAACCAGGTTTT |
| eIF2B4-1_spacer-rev | CTCTAAAACCTGGTTGTTGAATCTGGTCTC |

| | |
|---|---|
| eIF2B4-1_RT/PBS-fwd | GTGCGTCCCTCCGGTTGTTGAATCTG |
| eIF2B4-1_RT/PBS-rev | AAAACAGATTCAACAACCGGAGGGAC |
| eIF2B4-2_spacer-fwd | CACCGTCCCTGGAGAGTTCCTCACTGTTTT |
| eIF2B4-2_spacer-rev | CTCTAAAACAGTGAGGAACTCTCCAGGGAC |
| eIF2B4-2_RT/PBS-fwd | GTGCACAACACcgCCtAGTGAGGAACTCTCCA |
| eIF2B4-2_RT/PBS-rev | AAAATGGAGAGTTCCTCACTaGGcgGTGTTGT |
| eIF2B5_spacer-fwd | CACCGCCATCACCACGTTGTCCTCAGTTTT |
| eIF2B5_spacer-rev | CTCTAAAACTGAGGACAACGTGGTGATGGC |
| eIF2B5_RT/PBS-fwd | GTGCACACGCTGCCATGAGGACAACGT |
| eIF2B5_RT/PBS-rev | AAAAACGTTGTCCTCATGGCAGCGTGT |
| otc_spacer-fwd | CACCGACCACACAAGACATTCACTTGTTTT |
| otc_spacer-rev | CTCTAAAACAAGTGAATGTCTTGTGTGGTC |
| otc_RT/PBS-fwd | GTGCACCGAGCGGTGTCTGTGAGACTTTCATTCACACCCAAGTGAATGTCTTGTG |
| otc_RT/PBS-rev | AAAACACAAGACATTCACTTGGGTGTGAATGAAAGTCTCACAGACACCGCTCGGT |
| gabaR1α_spacer-fwd | CACCGCCACAGACTTCTTTCCATTGGTTTT |
| gabaR1α_spacer-rev | CTCTAAAACCAATGGAAAGAAGTCTGTGGC |
| gabaR1α_RT/PBS-fwd | GTGCATACATTTTTCCACAATGGAAAGAAG |
| gabaR1α_RT/PBS-rev | AAAACTTCTTTCCATTGTGGAAAAATGTAT |
| Dnmt1_spacer-fwd | caccGCGGGCTGGAGCTGTTCGCGCgtttt |
| Dnmt1_spacer-rev | ctctaaaacGCCTAATGTACTGTGTGCAGc |
| Dnmt1_RT/PBS-fwd | gtgcAAGATGgCAGCGCGAACAGCTCCAG |
| Dnmt1_RT/PBS-rev | aaaaCTGGAGCTGTTCGCGCTGCCATCTT |
| Dnmt1_sgRNA-fwd | caccGTCGTCTGCAACCTGCAAGA |
| Dnmt1_sgRNA-rev | aaacTCTTGCAGGTTGCAGACGAC |
| mPKU1_spacer-fwd | caccGCCTAATGTACTGTGTGCAGgtttt |
| mPKU1_spacer-rev | ctctaaaacCTGCACACAGTACATTAGGC |
| mPKU1.1_RT/PBS-fwd | gtgcTCCGaGTCTcCCACTGCACACAGTACATT |
| mPKU1.1_RT/PBS-rev | aaaaAATGTACTGTGTGCAGTGGgAGACtCGGA |
| mPKU1.2_RT/PBS-fwd | gtgcCCTTCCGaGTCTcCCACTGCACACAGTACATT |
| mPKU1.2_RT/PBS-rev | aaaaAATGTACTGTGTGCAGTGGgAGACtCGGAAGG |
| mPKU2_spacer-fwd | caccGCCTAATGTACTGTGTGCAGTgtttt |
| mPKU2_spacer-rev | ctctaaaacACTGCACACAGTACATTAGGC |
| mPKU2.1_RT/PBS-fwd | gtgcTTCCGgGTCTtCCACTGCACACAGTACAT |
| mPKU2.1_RT/PBS-rev | aaaaATGTACTGTGTGCAGTGGAAGACCCGGAA |
| mPKU2.2_RT/PBS-fwd | gtgcGCCTTCCGgGTCTtCCACTGCACACAGTACAT |
| mPKU2.2_RT/PBS-rev | aaaaATGTACTGTGTGCAGTGGAAGACCCGGAAGGC |
| mPKU3_spacer-fwd | caccgATGTACTGTGTGCAGTGGgAgtttt |
| mPKU3_spacer-rev | ctctaaaacTCCCACTGCACACAGTACATc |
| mPKU3.1_RT/PBS-fwd | gtgcGCCTTCCGAGTCTTcCACTGCACACAGTA |
| mPKU3.1_RT/PBS-rev | aaaaTACTGTGTGCAGTGGAAGACTCGGAAGGC |
| mPKU3.2_RT/PBS-fwd | gtgcGCCTGGCCTTCCGAGTCTTcCACTGCACACAGT |
| mPKU3.2_RT/PBS-rev | aaaaACTGTGTGCAGTGGAAGACTCGGAAGGCCAGGC |
| mPKU_sgRNA-fwd | caccgCTTGGGTGGCCTGGCCTTCC |
| mPKU_sgRNA-rev | aaacGGAAGGCCAGGCCACCCAAGc |
| mPKU_ngRNAPE3-2-fwd | caccCCTGGCCTTCCGAGTCTTtCC |
| mPKU_ngRNAPE3b-2-rev | aaacGGAAGACTCGGAAGGCCAGG |
| mPKU_ngRNAPE3b-3-fwd | caccGCCTGGCCTTCCGAGTCTTtC |
| mPKU_ngRNAPE3b-3-rev | aaacGAAGACTCGGAAGGCCAGGC |
| SpCas_scaffold-fwd | AGAGCTAGAAATAGCAAGTTAAAATAAGGCTAGTCCGTTATCAACTTGAAAAAGT GGCACCGAGTCG |
| SpCas_scaffold-rev | GCACCGACTCGGTGCCACTTTTTCAAGTTGATAACGGACTAGCCTTATTTTAACT TGCTATTTCTAG |
| rSTOP-Sa_spacer-fwd | CACCGTACTGAGGGGTACAATCCTACGTTTT |
| rSTOP-Sa_spacer-rev | TACTAAAACGTAGGATTGTACCCCTCAGTAC |
| rSTOP-Sa_RT/PBS-fwd | GAGAgGGACTAGAGTGGGATTGTACCCCT |
| rSTOP-Sa_RT/PBS-rev | AAAAAGGGGTACAATCCCACTCTAGTCCC |
| rSTOP-Saur_spacer-fwd | CACCGAGAgccaccatgGGACTAGAGGTTTT |
| rSTOP-Saur_spacer-rev | TACTAAAACCTCTAGTCCcatggtggcTCT |

| | |
|---|---|
| rSTOP-Saur_RT/PBS-fwd | GAGAGGGTACAATCCcACTCTAGTCCcatggtg |
| rSTOP-Saur_RT/PBS-rev | AAAAcaccatgGGACTAGAGTgGGATTGTACCC |
| SaCas_scaffold-fwd | AGTActctggaaacagaatctactaaaacaaggcaaaatgccgtgtttatctcgtcaacttgttggc |
| SaCas_scaffold-rev | TCTCgccaacaagttgacgagataaacacggcattttgccttgttttagtagattctgtttccaga |
| rSTOP-Cj_spacer-fwd | CACCgAgccaccatgGGACTAGAGTaGGTTTT |
| rSTOP-Cj_spacer-rev | gactaaaacCtACTCTAGTCCcatggtggcTC |
| rSTOP-Cj_RT/PBS-fwd | ccgcGAGGGGTACAATCCcACTCTAGTCCcatg |
| rSTOP-Cj_RT/PBS-rev | aaaacatgGGACTAGAGTgGGATTGTACCCCTC |
| CjCas_scaffold-fwd | agtccctgaaaagggactaaaataaagagtttgcgggactctgcggggttacaatcccctaaaa |
| CjCas_scaffold-rev | gcggttttaggggattgtaaccccgcagagtcccgcaaactctttattttagtcccttttcagg |
| HEK3_spacer-fwd | CACCgcacgtgctcagtctgggccccGTTTT |
| HEK3_spacer-rev | TACTAAAACggggcccagactgagcacgtgc |
| HEK3_RT/PBS-fwd | GAGAtgggtcaatccttggtgcccagactgagca |
| HEK3_RT/PBS-rev | AAAAtgctcagtctgggcaccaaggattgaccca |
| HEK4_spacer-fwd | CACCgcccactgtagtcacacagcacGTTTT |
| HEK4_spacer-rev | TACTAAAACgtgctgtgtgactacagtgggc |
| HEK4_RT/PBS-fwd | GAGAagcggagactctggtActgtgtgactacag |
| HEK4_RT/PBS-rev | AAAActgtagtcacacagTaccagagtctccgct |
| AAVS1_spacer-fwd | CACCGTTAGGCAGATTCCTTATCTGGGTTTT |
| AAVS1_spacer-rev | gactaaaacCCAGATAAGGAATCTGCCTAAC |
| AAVS1_RT/PBS-fwd | ccgcGGGGGTGTGTCACCgGATAAGGAATCT |
| AAVS1_RT/PBS-rev | aaaaAGATTCCTTATCcGGTGACACACCCCC |
| AAVS2_spacer-fwd | CACCGGAGTGTGACAGCCTGGGGCCCGTTTT |
| AAVS2_spacer-rev | gactaaaacGGGCCCCAGGCTGTCACACTCC |
| AAVS2_RT/PBS-fwd | ccgcACCTGTGTGCCTGGaCCCCAGGCTGTC |
| AAVS2_RT/PBS-rev | aaaaGACAGCCTGGGGtCCAGGCACACAGGT |
| PKU_amplicon-fwd | cgacatccctcagtaatgcca |
| PKU_amplicon-rev | gcagtggatcatggggacca |

**Table 3: Nucleotide sequences of HTS amplicons.**

| amplicon | amplicon sequence |
|---|---|
| adrb1 | CCAGCATTGAGACCCTGTGTGTCATCGCCCTGGACCGCTACCTCGCCATCACGTCGCCCTTTCGCTACCAGAGTTTGCTGACGCGCGCGCGAGCGCGGGCCCTCGTGTGCACAGTGTGGGCCATCTCGGCGTTGGTGTCCTTCCTGCCCATCCTCATG |
| app | TGGGTAGGCTTTGTCTTACAGTGTTATTATTTATGAGTAAAACTAATTGGTTGTCCTGCATACTTTAATTATGATGTAATACAGGTTCTGGGTTGACAAATATCAAGACGGAGGAGATCTCTGAAGTGAAGATGGATgCAGAATTCCGACATGACTCAGGATAT |
| eIF2B4-1 | GTGCTGGAATTCCTAAAAGGATGTTGACACCCACTTGTCATTTCAGATGATCCCGATGATCTGCAGTGTAAGCGGGGAGACCAGGTGGCCCTGGCTAACTGGCAGAGCCACCCGTCCCTCTGGTTGTTGAATCTGGTCTATGACGTGACTCCACCTGAGCTCGTGGATCTGGTGATCAC |
| eIF2B4-2 | GGATCCACTAGTAACGGCCGCCAGTGTGCTGGAATTCGAAGCAATGACTCTCACAATCTGAACACTTACTTCCCTAGTGAAGCTTACCCCTGTCCCATAGTAGAGATCTTCCTTAAGGGGTGTTTTGTATTCTCTAGGTGATTCAGGATTACACAACACtgCCtAGTGAGGAACTCTCCAGGGATCTTGTAAATAAACTAAAACCCTACATCAGGTGAGCACAGCC |
| eIF2B5 | GATCACAGAGTTGGGCAGACTAACTGTGCCTCTGGTTCTTAATAGGTTAAGAAGGAAGCTAGAAAAAAATGTCTCTGTGATGACAATGGTCTTCAAAGAGTCGTCACCCAGCCACCCTACACACTGCCATGAGGACAACGTGGTGATGGCTGTGGACAGCGCCACCAACAGGGTTCTTCACTTCCAGAAGACCCAAGG |
| otc | GGGAGGACACCCTTCCTTTCTTACCACACAAGACATTCACTTGGGTGTGAATGAAAGTCTCACAGACACCGCTCAGTTTGTAAAACTTTTCTTCCTTCCAAAGTTTATTTCAAACTCTGATGGGTTAGTTTAAAAGAGAAGATGATGCTTCTCCTTAGATAATGGTCTCCCCGGGTGGGCACAGGTAGGGACTG |
| gabaR1α | CGTCAAAGTTGGAAGGATGAAAGATTAAAATTCAAAGGACCCATGACAGTGCTCCGGCTAAACAACCTTATGGCCAGTAAAATCTGGACTCCAGATACATTTTTCCGCAATGGAAAGAAGTCTGTGGCCCACAACATGACCATGCCTAATAAGCTCCTGCGTATCACAGAGGATGGCGGTCCTATGAATA |

| | |
|---|---|
| Dnmt1 | GTCTTCCCCCACTCTCTTGCCCTGTGTGGTACATGCTGCTTCCGCTTGCGCCGCCCCCTCCCAATTG<br>GTTTCCGCGCGCGCGAAAAAGCCGGGGTCTCGTTCAGAGCTGTTCTGTCGTCTGCAACCTGCAAGAT<br>GCCAGCGCGAACAGCTCCAGCCCGAGTGCCTGCGCTTGCCTCCCCGGCAGGCTCGCTCCCGGACCAT<br>GTCCGCAGGCGGTAGGTGCCACGCAGGGTGGGGGTGAGGGGCGGGACCGATGCCGAGGCATATATTG<br>GGGG |
| PKU | CCGTCCTGTTGCTGGCTTACTGTCGTCTCGAGATTTCTTGGGTGGCCTGGCCTTCCGaGTCTcCCAC<br>TGCACACAGTACATTAGGCATGGATCTAAGCCCATGTACACACCTGAACCGTAAGTATCATTCTTCA<br>GCTACCCCTGCCAACCACAATGGATGCTCA |
| HEK3 | agacagggatcccagggaaacgcccatgcaattagtctatttctgctgcaagtaagcatgcatttgt<br>aggcttgatgctttttttctgcttctccagccctggcctgggtcaatccttggggcccagactgagc<br>acgtgatggcagaggaaaggaagccctgcttcctccagagggcgtcgcaggacagcttttcctagac<br>aggggctagtatgtgcagctcctgcaccgggatactggttgacaagtttggctggg |
| HEK4 | ccctcccttcaagatggctgacaaaggccgggctgggtggaaggaagggaggaagggcgaggcagag<br>ggtccaaagcaggatgacaggcaggggcaccgcggcgccccggtggcactgcggctggaggtggggg<br>ttaaagcggagactctggtgctgtgtgactacagtgggggccctgccctctctgagcccccgcctcc<br>aggcctgtgtgtgtgtctccgtt |
| AAVS1 | CCTTAGGCCTCCTCCTTCCTAGTCTCCTGATATTGGGTCTAACCCCCACCTCCTGTTAGGCAGATTC<br>CTTATCTGGTGACACACCCCCATTTCCTGGAGCCATCTCTCTCCTTGCCAGAACCTCTAAGGTTTGC<br>TTACGATGGAGCCAGAGAGGAT |
| AAVS2 | AGAATGAGAAACGGTGGCCCGTGTCAGCCCCTGGCTGCAGGGCCCCGTGCAGAGGGGGCCTCAGTGA<br>ACTGGAGTGTGACAGCCTGGGGCCCAGGCACACAGGTGTGCAGCTGTCTCACCCCTCTGGGAGTCCC<br>GCCCAGGCCCCTGAGTCTGTCCCAGCACAGGGTGGCCTTCCTCCACCCTGCATAGC |
| OT1-peg | GGTTTCCCATTTCCCATCATCATTCACCTGTCCTTCCCACTGCACACAGTACATGCTCCG<br>TAGCCCAACAGTCCTGCAGGGCCCATACATGTACACACTTCCCAG |
| OT2-peg | AGCATGTATGGGTTCCGAGGCTGGGCATGAGCAGCCTCCTGGAATTTCATGGACCTTGTC<br>CTGCACACAGTGCATTAGGCAAGGTCTTGCAACAAGGCCTTATTTACTAGTGTTACCAGG<br>ACCGTGCTGAAACTGGTTTC |
| OT3-peg | ATTAGGGAGGAGGGTAGAAGTGTTGGTAACCACAGGGTGGGTTGGAACCACTGCCATATT<br>TAATGTACTGTGTGGTAGGCAGCGGAAGCTCATGTACTGC |
| OT4-peg | CAGGGCAAGCAGGTAGATGTTCCTTCGCAGCTTCCCACTGCAGAGTAATGTACTCTGGGC<br>AGTGGGTGTGCTCCAGCTCAGCTTGCCCAAACTGAAGGGCGACAGGT |
| OT5-peg | AGTATACCACTGTTTGTGCATTCCTCTTGTGTCTGTGCCTTGGTGCACACATATGTACTG<br>TGTGCAGAGGTATGTATTCTATTGTGTGCAGCGTACTTGC |

**Table 4: Oligos used for RT-qPCR.**

| oligo name | oligo sequence |
|---|---|
| SYBR_pegRNA_Fw | GTTTTAGAGCTAGAAATAGC |
| SYBR_pegRNA_Rev | GCACCGACTCGGTGCCACTT |
| SYBR_N-int_Fw | CAGTGGCATGATAGAGGCGA |
| SYBR_N-int_Rev | GCATTTGTCCGTCAACCGTC |
| SYBR_RT_Fw | CCCCTTGCCTACGGATGGTA |
| SYBR_RT_Rev | TGTCCAAAAGCAAGGCCTGATA |
| SYBR_Cas_Fw | TGATCGAGACAAACGGCGAA |
| SYBR_Cas_Rev | CTGTCTGCACCTCGGTCTTT |
| SYBR_mRPLP0_FW | TGAGATTCGGGATATGCTGTTGG |
| SYBR_mRPLP0_Rev | CGGGTCCTAGACCAGTGTTCT |

**Table 5: Amino acid sequences of PE constructs.**

| p.NLS-Cas9(H840A)-xten-RT-NLS: |
|---|
| MKRTADGSEFESPKKKRKVDKKYSIGLDIGTNSVGWAVITDEYKVPSKKFKVLGNTDRHSIKKNLIGALLFDSG<br>ETAEATRLKRTARRRYTRRKNRICYLQEIFSNEMAKVDDSFFHRLEESFLVEEDKKHERHPIFGNIVDEVAYHEK<br>YPTIYHLRKKLVDSTDKADLRLIYLALAHMIKFRGHFLIEGDLNPDNSDVDKLFIQLVQTYNQLFEENPINASGV<br>DAKAILSARLSKSRRLENLIAQLPGEKKNGLFGNLIALSLGLTPNFKSNFDLAEDAKLQLSKDTYDDDLDNLLAQ<br>IGDQYADLFLAAKNLSDAILLSDILRVNTEITKAPLSASMIKRYDEHHQDLTLLKALVRQQLPEKYKEIFFDQSKN<br>GYAGYIDGGASQEEFYKFIKPILEKMDGTEELLVKLNREDLLRKQRTFDNGSIPHQIHLGELHAILRRQEDFYPFL<br>KDNREKIEKILTFRIPYYVGPLARGNSRFAWMTRKSEETITPWNFEEVVDKGASAQSFIERMTNFDKNLPNEKVL<br>PKHSLLYEYFTVYNELTKVKYVTEGMRKPAFLSGEQKKAIVDLLFKTNRKVTVKQLKEDYFKKIECFDSVEISG |

VEDRFNASLGTYHDLLKIIKDKDFLDNEENEDILEDIVLTLTLFEDREMIEERLKTYAHLFDDKVMKQLKRRRYT
GWGRLSRKLINGIRDKQSGKTILDFLKSDGFANRNFMQLIHDDSLTFKEDIQKAQVSGQGDSLHEHIANLAGSPA
IKKGILQTVKVVDELVKVMGRHKPENIVIEMARENQTTQKGQKNSRERMKRIEEGIKELGSQILKEHPVENTQLQ
NEKLYLYYLQNGRDMYVDQELDINRLSDYDVDAIVPQSFLKDDSIDNKVLTRSDKNRGKSDNVPSEEVVKKMK
NYWRQLLNAKLITQRKFDNLTKAERGGLSELDKAGFIKRQLVETRQITKHVAQILDSRMNTKYDENDKLIREVK
VITLKSKLVSDFRKDFQFYKVREINNYHHAHDAYLNAVVGTALIKKYPKLESEFVYGDYKVYDVRKMIAKSEQ
EIGKATAKYFFYSNIMNFFKTEITLANGEIRKRPLIETNGETGEIVWDKGRDFATVRKVLSMPQVNIVKKTEVQT
GGFSKESILPKRNSDKLIARKKDWDPKKYGGFDSPTVAYSVLVVAKVEKGKSKKLKSVKELLGITIMERSSFEK
NPIDFLEAKGYKEVKKDLIIKLPKYSLFELENGRKRMLASAGELQKGNELALPSKYVNFLYLASHYEKLKGSPED
NEQKQLFVEQHKHYLDEIIEQISEFSKRVILADANLDKVLSAYNKHRDKPIREQAENIIHLFTLTNLGAPAAFKYF
DTTIDRKRYTSTKEVLDATLIHQSITGLYETRIDLSQLGGDSGGSSGGSSGSETPGTSESATPESSGGSSGGSSTLNI
EDEYRLHETSKEPDVSLGSTWLSDFPQAWAETGGMGLAVRQAPLIIPLKATSTPVSIKQYPMSQEARLGIKPHIQ
RLLDQGILVPCQSPWNTPLLPVKKPGTNDYRPVQDLREVNKRVEDIHPTVPNPYNLLSGLPPSHQWYTVLDLKD
AFFCLRLHPTSQPLFAFEWRDPEMGISGQLTWTRLPQGFKNSPTLFNEALHRDLADFRIQHPDLILLQYVDDLLL
AATSELDCQQGTRALLQTLGNLGYRASAKKAQICQKQVKYLGYLLKEGQRWLTEARKETVMGQPTPKTPRQL
REFLGKAGFCRLFIPGFAEMAAPLYPLTKPGTLFNWGPDQQKAYQEIKQALLTAPALGLPDLTKPFELFVDEKQ
GYAKGVLTQKLGPWRRPVAYLSKKLDPVAAGWPPCLRMVAAIAVLTKDAGKLTMGQPLVILAPHAVEALVKQ
PPDRWLSNARMTHYQALLLDTDRVQFGPVVALNPATLLPLPEEGLQHNCLDILAEAHGTRPDLTDQPLPDADHT
WYTDGSSLLQEGQRKAGAAVTTETEVIWAKALPAGTSAQRAELIALTQALKMAEGKKLNVYTDSRYAFATAHI
HGEIYRRRGWLTSEGKEIKNKDEILALLKALFLPKRLSIIHCPGHQKGHSAEARGNRMADQAARKAAITETPDTS
TLLIENSSPSGGSKRTADGSEFEPKKKRKV*

p.NLS-Cas9(H840A)-xten-RTΔ*RnH*-NLS:

MKRTADGSEFESPKKKRKVDKKYSIGLDIGTNSVGWAVITaaDEYKVPSKKFKVLGNTDRHSIKKNLIGALLFDS
GETAEATRLKRTARRRYTRRKNRICYLQEIFSNEMAKVDDSFFHRLEESFLVEEDKKHERHPIFGNIVDEVAYHE
KYPTIYHLRKKLVDSTDKADLRLIYLALAHMIKFRGHFLIEGDLNPDNSDVDKLFIQLVQTYNQLFEENPINASG
VDAKAILSARLSKSRRLENLIAQLPGEKKNGLFGNLIALSLGLTPNFKSNFDLAEDAKLQLSKDTYDDDLDNLLA
QIGDQYADLFLAAKNLSDAILLSDILRVNTEITKAPLSASMIKRYDEHHQDLTLLKALVRQQLPEKYKEIFFDQSK
NGYAGYIDGGASQEEFYKFIKPILEKMDGTEELLVKLNREDLLRKQRTFDNGSIPHQIHLGELHAILRRQEDFYPF
LKDNREKIEKILTFRIPYYVGPLARGNSRFAWMTRKSEETITPWNFEEVVDKGASAQSFIERMTNFDKNLPNEKV
LPKHSLLYEYFTVYNELTKVKYVTEGMRKPAFLSGEQKKAIVDLLFKTNRKVTVKQLKEDYFKKIECFDSVEIS
GVEDRFNASLGTYHDLLKIIKDKDFLDNEENEDILEDIVLTLTLFEDREMIEERLKTYAHLFDDKVMKQLKRRRY
TGWGRLSRKLINGIRDKQSGKTILDFLKSDGFANRNFMQLIHDDSLTFKEDIQKAQVSGQGDSLHEHIANLAGSP
AIKKGILQTVKVVDELVKVMGRHKPENIVIEMARENQTTQKGQKNSRERMKRIEEGIKELGSQILKEHPVENTQL
QNEKLYLYYLQNGRDMYVDQELDINRLSDYDVDAIVPQSFLKDDSIDNKVLTRSDKNRGKSDNVPSEEVVKKM
KNYWRQLLNAKLITQRKFDNLTKAERGGLSELDKAGFIKRQLVETRQITKHVAQILDSRMNTKYDENDKLIREV
KVITLKSKLVSDFRKDFQFYKVREINNYHHAHDAYLNAVVGTALIKKYPKLESEFVYGDYKVYDVRKMIAKSE
QEIGKATAKYFFYSNIMNFFKTEITLANGEIRKRPLIETNGETGEIVWDKGRDFATVRKVLSMPQVNIVKKTEVQ
TGGFSKESILPKRNSDKLIARKKDWDPKKYGGFDSPTVAYSVLVVAKVEKGKSKKLKSVKELLGITIMERSSFEK
NPIDFLEAKGYKEVKKDLIIKLPKYSLFELENGRKRMLASAGELQKGNELALPSKYVNFLYLASHYEKLKGSPED
NEQKQLFVEQHKHYLDEIIEQISEFSKRVILADANLDKVLSAYNKHRDKPIREQAENIIHLFTLTNLGAPAAFKYF
DTTIDRKRYTSTKEVLDATLIHQSITGLYETRIDLSQLGGDSGGSSGGSSGSETPGTSESATPESSGGSSGGSSTLNI
EDEYRLHETSKEPDVSLGSTWLSDFPQAWAETGGMGLAVRQAPLIIPLKATSTPVSIKQYPMSQEARLGIKPHIQ
RLLDQGILVPCQSPWNTPLLPVKKPGTNDYRPVQDLREVNKRVEDIHPTVPNPYNLLSGLPPSHQWYTVLDLKD
AFFCLRLHPTSQPLFAFEWRDPEMGISGQLTWTRLPQGFKNSPTLFNEALHRDLADFRIQHPDLILLQYVDDLLL
AATSELDCQQGTRALLQTLGNLGYRASAKKAQICQKQVKYLGYLLKEGQRWLTEARKETVMGQPTPKTPRQL
REFLGKAGFCRLFIPGFAEMAAPLYPLTKPGTLFNWGPDQQKAYQEIKQALLTAPALGLPDLTKPFELFVDEKQ
GYAKGVLTQKLGPWRRPVAYLSKKLDPVAAGWPPCLRMVAAIAVLTKDAGKLTMGQPLVILAPHAVEALVKQ
PPDRWLSNARMTHYQALLLDTDRVQFGPVVALNPATLLPGSEFEPKKKRKV*

**Supplementary Data 1.** Complete images of Western Blots.

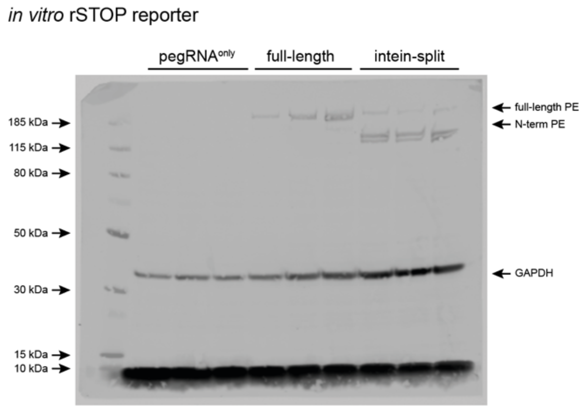


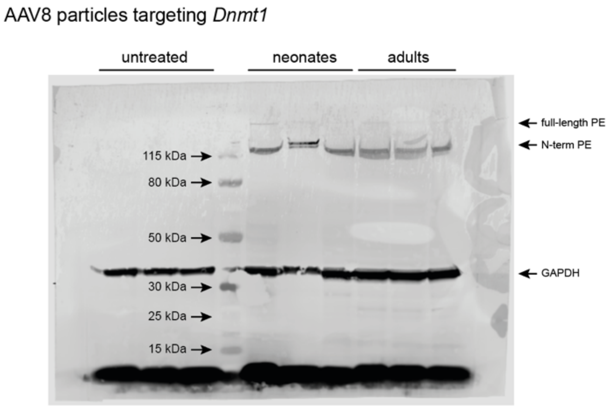
